# Tricuspid Valve Chordae Tendineae: Where are They, How Many are There, and What are Their Properties?

**DOI:** 10.64898/2026.11.10.765710

**Authors:** Collin E. Haese, Colton J. Kostelnik, Vijay K. Dubey, Sumedh Seetharam, Mrudang Mathur, Davide Tondi, Matthew M. Duda, Kelly C. Higa, Robyn Fong, Marcin Malinowski, Magda Piekarska, Adrian Buganza Tepole, Jan N. Fuhg, Issam D. Moussa, Matthew R. Summers, Emiliano Votta, William Hiesinger, Tomasz A. Timek, Manuel K. Rausch

**Affiliations:** Walker Department of Mechanical Engineering, The University of Texas at Austin, 204 E Dean Keeton, Austin, TX, 78712, United States of America; Department of Biomedical Engineering, The University of Texas at Austin, 107 W Dean Keeton St, Austin, TX, 78712, United States of America; Department of Cardiothoracic Surgery, Stanford University, 870 Quarry Rd Extension, Palo Alto, CA, 94305, United States of America; Department of Electronics, Information and Bioengineering, Politecnico di Milano, 34 Via Giuseppe Ponzio, Milan, Italy; Division of Cardiothoracic Surgery, Corewell Health, 100 Michigan Ave SE, Grand Rapids, MI, 49503, United States of America; Department of Cardiac Surgery, Wojewódzki Szpital im. Sw. Ojca Pio, Przemyśl, Poland; Department of Mechanical Engineering, Columbia University, 500 W 120th St, New York, NY, 10027, United States of America; Department of Aerospace Engineering & Engineering Mechanics, The University of Texas at Austin, 2617 Wichita St, Austin, TX, 78712, United States of America; Oden Institute for Computational Engineering and Sciences, The University of Texas at Austin, 201 E 24th St, Austin, TX, 78712, United States of America; Carle Illinois College of Medicine, University of Illinois at Urbana-Champaign, 506 S Mathews Ave, Urbana, IL, 61801, United States of America; Sentara Heart Valve Center, Sentara Heart Hospital, 600 Gresham Dr, Norfolk, VA, 23507, United States of America; College of Human Medicine, Michigan State University, 15 Michigan St NE, Grand Rapids, MI, 49503, United States of America

**Keywords:** Tissue Mechanics, Constitutive Modeling, Hierarchical Bayesian Modeling, Open Data, Mechanical Testing

## Abstract

**Background:** Tricuspid chordae play an important role in valve function. Unfortunately, they are poorly resolved in clinical images and their mechanical properties cannot be measured noninvasively. Limited human anatomical and mechanical data hinder our understanding of the tricuspid valve in general and efforts to model the valve in particular.

**Objectives:** Our primary objective was to quantify chordal insertion patterns, structure, and mechanical properties. Our secondary objective was to test whether chordal structure and properties differed between control hearts and hearts with heart failure.

**Methods:** We mapped chordal insertions in 25 excised human valves, measured the cross-sectional areas of 281 specimens, and conducted uniaxial tensile tests on 265 specimens from 12 of the 25 valves (eight control and four heart failure). We used mixed-effects models to assess the influence of heart failure and anatomical position. Finally, we used a hierarchical Bayesian model to represent area-dependent Ogden material parameters.

**Results:** We found that chordal insertion rates were highest at the free edge (p *<* 0.001), and lowest in the anterior leaflet (p *<* 0.001). Median cross-sectional area was 0.223 mm^2^, with anterior chordae the largest (p *<* 0.001) and septal the smallest (p *<* 0.001). A 10% increase in cross-sectional area was associated with 5.3% and 3.3% decreases in toe and calf stiffness, respectively. Heart failure was not associated with an overall increase in chordal area or material stiffness.

**Conclusions:** Human chordal geometry and material behavior varied with location and size. These openly shared data support assigning chordal insertions, sizes, and material properties in tricuspid valve models.

## 1. Introduction

The tricuspid valve (TV) ensures unidirectional blood flow from the right atrium into the right ventricle. Its function depends on the coordinated interaction between the annulus, leaflets, and chordae tendineae [1]. The latter connect the leaflets to the papillary muscles, direct leaflet motion, prevent prolapse, and distribute forces during closure. Disruption of this coordination can cause incomplete coaptation and backflow of blood, i.e., regurgitation. Tricuspid regurgitation (TR) is a common condition [2] and, when severe, it is associated with increased morbidity and mortality [3]. Many repair and replacement techniques address TR, but opportunities remain to improve and optimize these strategies [4, 5, 6, 7]. Computational models provide a controlled and repeatable means to systematically investigate repair techniques and evaluate how geometry, tissue properties, and repair strategies affect valve mechanics [8, 9, 10]. The accuracy of these models, however, depends on the geometric and mechanical data used to represent the valve [11].

For models that represent the entire valvular apparatus, these inputs include the location, size, and mechanical behavior of the chordae tendineae. Human tricuspid chordae, however, remain insufficiently characterized [12, 13]. Anatomical studies of the human TV have described chordal classifications and distributions [14, 15, 16], but mechanical evidence is derived largely from mitral [17, 18], porcine [13, 19], or ovine tricuspid chordae tendineae [20]. Human mechanical data remain limited to small studies by Lim and colleagues [21, 22], measurements reported in our previous computational studies [23, 24], and one dissertation chapter examining fresh human chordae [25]. Together, these studies provide an initial basis for understanding human tricuspid chordal mechanics but offer limited information about how mechanical properties vary between individual valves and by anatomical position.

Beyond positional differences, chordal properties may also vary in response to changes in the mechanical and hemodynamic environment. Previous studies have demonstrated that tricuspid valve leaflets undergo remodeling in response to altered loading conditions, including tissue thickening and stiffening in heart failure models [26, 27, 28]. Similarly, changes in the number of mitral chordal attachments have been observed during pregnancy [29]. These findings suggest that tricuspid chordae may similarly thicken and stiffen in response to altered loading conditions associated with heart failure [30, 31]. However, whether human tricuspid chordal size and material properties differ between control hearts and hearts with heart failure remains unknown.

The limited characterization of human tricuspid chordal anatomy and mechanics also presents challenges for image-based computational modeling. High-fidelity subject-specific models have been constructed using images and mechanical measurements of excised valves [23, 24]. However, routine clinical use of computational models will require building them without access to excised tissue. One critical issue in building these models is that the subvalvular apparatus, particularly the chordae tendineae, is often incompletely resolved in clinical images [32, 33]. Image-based models must supplement the available geometry with assumptions about chordal anatomy and mechanics. Recent approaches have constructed synthetic chordal architectures or represented chordal support through equivalent loads or displacements [34, 35, 36]. Synthetic architectures can produce functionally equivalent models [37, 38, 39], although model behavior is sensitive to the number and location of chordal insertions [37, 11]. Furthermore, these techniques often assume simplified chordal insertion patterns that may not be anatomically realistic. Thus, while they can reliably reproduce the imaged valve configuration, they may fail to predict the valve’s response to interventions that alter subvalvular geometry.

Beyond insertion patterns, chordal size and material properties also determine the mechanical response of models that explicitly represent the chordae. Because comprehensive human data are unavailable, existing models commonly use material properties measured in individual excised valves [23, 40] or extrapolate from mitral [41, 42] and animal tissues [43, 44]. Thus, comprehensive measurements of human chordal insertion patterns, sizes, and mechanical behavior are needed to assign chordal architectures and material properties when direct measurements are unavailable from clinical images.

To address these gaps, our primary objective was to characterize human tricuspid chordae by answering three questions: 1) Where and how many chordae tendineae insert into each leaflet? 2) How large are these chordae? 3) What are their mechanical properties? Our secondary objective was to test the hypothesis that chordae from hearts with heart failure would have larger cross-sectional areas and higher material stiffness than those from control hearts, consistent with remodeling in an altered mechanical and hemodynamic environment. To this end, we mapped chordal insertions in excised tricuspid valves, measured chordal cross-sectional areas, performed uniaxial tensile testing, and evaluated differences by heart-failure status and anatomical location.

## 2. Methods

### 2.1. Collection and Excision

This study was deemed non-human subjects research by Corewell Health, UT-Austin, and Stanford University Institutional Review Boards. We collected 25 human hearts from 2017–2026. Fourteen hearts were rejected from transplantation due to mild to severe left-sided disease or infection risk but had normal right ventricular function and presented with no more than trace TR. We classified these as “control” (CTL). The remaining eleven hearts were obtained from patients undergoing transplantation for heart failure (HF). Subject data are provided in Supplementary Table S1. We excised each TV, cut it at the posterior-septal commissure, laid the valve flat on a calibrated mat, and photographed the ventricular surface of the leaflets (Figure 1A). Subsequently, we isolated the leaflets and cooled them to –80 ^°^C in a cryogenic solution of 9:1 DMEM to DMSO with Pierce protease inhibitor mini tablets (Thermo Scientific, A32953, Waltham, MA, USA) at a rate of –1^°^C/min using a Mr. Frosty container (Thermo Scientific, 5100-0001)[26].

**Figure 1:**
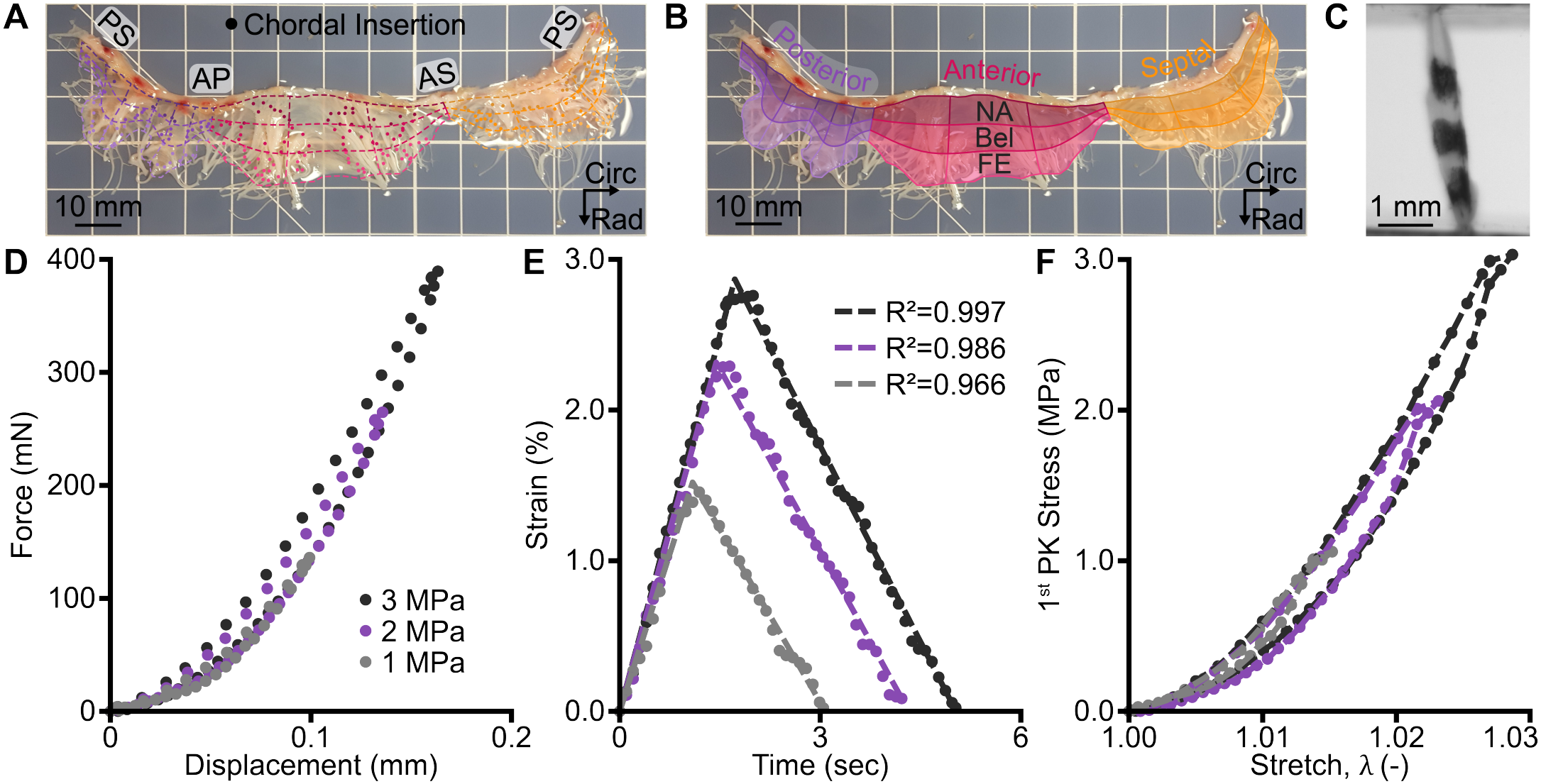
Characterization of human tricuspid valve chordae tendineae. **(A)** We marked all chordal insertions on images in a 3×3 radial-by-circumferential grid defined across each leaflet scallop. **(B)** We excised chordae, recorded their leaflet and radial insertion site, and measured their cross-sectional area. We mechanically tested the chordae by **(C)** marking three points along each chorda and **(D)** recording the uniaxial force-displacement response required to attain 1, 2, or 3 MPa peak 1^*st*^ Piola-Kirchhoff stress. **(E)** We measured nominal strain by tracking the markers and fit linear functions to the strain-time data of the final loading-unloading cycles to 1, 2, or 3 MPa peak 1^*st*^ Piola-Kirchhoff stress. **(F)** Lastly, we converted the data into 1^st^ Piola-Kirchhoff stress versus stretch (*λ*). Panels **(D-F)** show data from one specimen.

### 2.2. Density Mapping

To quantify the spatial distribution of the chordae, we manually marked all identifiable chordal insertions on images of the ventricular surface of each valve. Using a custom MATLAB script (MATLAB 2024b, Mathworks, Natick, MA), we corrected perspective distortion and traced the annulus and the free edge. We visually identified and marked each chordal insertion site where the chordae tendineae terminated on the leaflet. In cases where a chorda bifurcated before reaching the leaflet, each of the terminal branches was marked. We divided each leaflet into scallops determined by indentations in the free edge and delineated scallops using line segments connecting the annulus to the free edge. We used a bilinear blended Coons patch [45], followed by Winslow smoothing [46, 47], to parametrize each scallop[48]. We divided each scallop into nine cells organized in a 3×3 mesh, with three circumferential regions aligned along the annulus, and three radial regions extending from the annulus-to-free edge (Figure 1A). We labeled the left (L), central (C), and right (R) circumferential regions as viewed from the ventricular surface, and the radial regions near-annulus (NA), belly (Bel), and free edge (FE). We measured cell areas using the calibrated length of each mat and calculated insertion density within each cell by dividing the insertion count by its area, with cells containing no insertions retained as zero-density observations.

### 2.3. Mechanical Testing

For mechanical testing, we thawed tissue from 12 valves (8 CTL, 4 HF) and excised the largest chordae from each leaflet region. We trimmed away any branches to isolate the main segment. Following our prior work [20], we classified chordae specimens as inserting into the septal (Sept), anterior (Ant), or posterior (Post) leaflet, at either the near-annulus (NA), belly (Bel), or free edge (FE) radial position (Figure 1B). Circumferential position was not recorded. Next, we measured each specimen’s diameter at three locations using a 4× objective on a light microscope (BX53 Upright Microscope, Olympus, Tokyo, Japan). We calculated the cross-sectional area using the mean diameter and assumed it to be circular. We further classified specimens based on their cross-sectional area as small (Sml; *<* 0.159 mm^2^), medium (Med; 0.159 − 0.315 mm^2^), or large (Lrg; *>* 0.315 mm^2^), which yielded approximately equal group sizes. In total we collected 281 chordae tendineae, see Table 1. Eleven specimens from one valve (CTL 3) lacked insertion site information and therefore were excluded from analyses involving that classification. Supplementary Table S3 provides specimen marginal and pairwise counts.

**Table 1:** Number of chordae tendineae by heart failure status, leaflet, and insertion site. Insertion site was unavailable for eleven control chordae from one valve, so insertion-site counts sum to 270.

|  | **Leaflet** |  |  |  | **Insertion Site** |  |  |  |
|---|---|---|---|---|---|---|---|---|
|  | Posterior | Anterior | Septal | **Total** | Near-Annulus | Belly | Free Edge | **Total** |
| Control | 48 | 58 | 54 | **160** | 32 | 61 | 56 | **149** |
| Heart Failure | 37 | 42 | 42 | **121** | 29 | 43 | 49 | **121** |
| **Total** | **85** | **100** | **96** | **281** | **61** | **104** | **105** | **270** |

We applied three fiducial markers along each specimen using a pigment liner pen (GEOTEC School and Office Supplies GmbH, Wörgl, Austria) as shown in Figure 1C. Then, we mounted specimens in custom stainless-steel clamps on a uniaxial testing device and removed slack. We used the cross-sectional area of each specimen to compute the force necessary to achieve 1, 2, and 3 MPa 1^st^ Piola-Kirchhoff stress. We preconditioned the specimen for 10 cycles at 0.1 mm/s to 1 MPa and recorded the axial force, clamp displacement, and images of the fiducial markers at 5 Hz during the final cycle, where the final unloading rate was 0.05 mm/s. With the specimen still mounted, we then repeated this testing method at 2 and 3 MPa. Specimens underwent 10 cycles at each sequential loading stage, without rest between stages, and preload was not recorded. An example of the force-displacement data collected for one specimen is shown in Figure **1D**. Specimens from three valves (CTL 3-5) were tested using a Cellscale UniVert with a 4.5 N load cell (accuracy of ± 9 mN; Cellscale, Waterloo, Ontario, Canada), recording at 5 Hz. For these specimens only, after the 3 MPa test, these specimens were loaded until failure. These data previously informed our subject-specific finite element models of the TV [23, 24]. Remaining specimens were tested to 1, 2, and 3 MPa using an Instron 5943 universal testing machine with a 10 N load cell (accuracy of ± 0.5% of reading; 2530-10N, Instron, Norwood, MA, USA), recording at 10 Hz. All tests were conducted in 1× PBS at 37 ^°^C within 8 hours of thawing. Due to specimen tearing, mounting difficulties, slippage, or other testing problems, 16 specimens were excluded from these mechanical analyses.

### 2.4. Data Processing

After mechanical testing, we processed these data using a custom MATLAB script. We calculated local axial strain via optical point tracking of the fiducial markers using LabJoy 11.18 (CellScale), where we averaged strain between marker pairs. For each test, the first image of the final loading cycle defined the strain reference, and the first force measurement of the final cycle was subtracted from all subsequent readings. Force and strain references were therefore reset at each final loading stage. To suppress high-frequency noise, we fit separate linear functions to the strain-time curves of the final loading and unloading strokes as shown in Figure **1E**. We enforced zero axial strain at the start of the final loading cycle and continuity in the strains between strokes.

Our linear strain-time fits had a median R^2^ of 0.968 [mean 0.955 ± 0.041 standard deviation, minimum: 0.807]. To account for recording rate differences, we linearly interpolated strain to the force-displacement recording time points. Lastly, we converted our force-displacement curves of the final cycle into 1^*st*^ Piola-Kirchhoff stress-stretch curves (Figure **1F**). Following our prior work [20], we calculated the toe stiffness, calf stiffness, transition stretch, and hysteresis for each test. Toe and calf stiffness were defined as the tangent slopes of the stress-stretch curve near 0 and 1 MPa, respectively (see Supplement S5.3). Transition stretch was defined as the stretch at which the tangent lines approximating the toe and calf stiffness intersected.

### 2.5. Constitutive Modeling

We fit ten constitutive models to the loading stroke of the stress-stretch curves. This included the eight hyperelastic models considered in our prior work [20], a three-term Ogden model [49], and a one-dimensional small-strain linear-elastic model. We assumed incompressible uniaxial deformation, with axial stretch *λ*_1_ = *λ* and off-axis stretches *λ*_2_ = *λ*_3_ = *λ*^−1*/*2^. Each constitutive model was defined by a strain-energy density function *W*(*λ*), from which the axial component of the 1^st^ Piola-Kirchhoff (PK) stress tensor, *P*_11_, was calculated as,

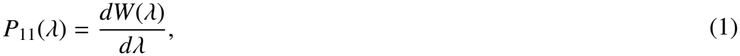

The uniaxial strain-energy functions and parameter bounds for all models are provided in Supplementary Table S4. For an Ogden model with N-terms, the 1^st^ PK axial stress is,

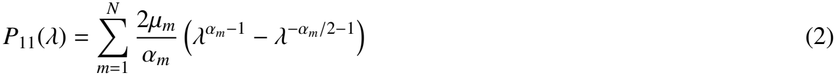

where *µ*_*m*_ has units of MPa and *α*_*m*_ is dimensionless.

We estimated model parameters by minimizing the error between the experimentally measured stress and the stress computed from each model using bounded nonlinear least-squares regression (`lsqnonlin`, MATLAB) with 20 multi-start initializations. For each model, we estimated a separate least-squares parameter vector 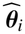 for each specimen *i* by fitting its available loading curves jointly. After fitting, we summarized specimen-level fit quality using pooled root mean square error (RMSE) and normalized mean square error goodness of fit (NMSE), calculated from unweighted stress residuals across the separate specimen-level fits.

In separate fits, we then estimated one parameter vector 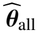 from the overall median response across all specimens and a separate vector 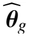 from the median response within each group *g* over 0–1 MPa. That is, we estimated one parameter vector per median curve. Groups were defined by heart failure, leaflet, insertion site, size, and leaflet-insertion site. We evaluated group fit quality with respect to the median curve using root mean square error (RMSE) and normalized mean square error goodness of fit (NMSE). The fitting objectives, construction of median curves, and error metrics are detailed in Supplement S3.

### 2.6. Bayesian Modeling

We fit a single hierarchical Bayesian model to estimate the one-term Ogden parameters, *µ*_1_ and *α*_1_, from the stress-stretch data following previously established approaches for Bayesian constitutive parameter identification in biological tissues [50, 51]. We modeled each log-transformed parameter as a linear function of the log-transformed chordal cross-sectional area, the sole fixed-effect covariate, with valve and specimen-within-valve included as random effects. The log parameterization ensured that *µ*_1_ and *α*_1_ were strictly positive, but no other explicit stability constraint was imposed during posterior sampling.

After tuning and fitting our model, we obtained a representative population-level response for each leaflet-insertion-site group based on that group’s median cross-sectional area. Here, we set valve and specimen random effects to zero. We log-transformed the median cross-sectional area for each group and standardized this area using the measured area distribution, then passed this through the fitted posterior of the Bayesian model. Then, we generated joint posterior draws of *µ*_1_ and *α*_1_ and selected a representative parameter pair from the joint draw whose predicted stress-stretch curve was closest to the pointwise posterior median curve. Thus, our Bayesian group predictions evaluate the same hierarchical fit at different cross-sectional areas. We also calculated pointwise 95% posterior intervals at each group’s median cross-sectional area. These intervals describe uncertainty in the population-level material response and do not account for variation among new valves or specimens, uncertainty in area, or observation noise. Further details are provided in Supplement S4.

### 2.7. Statistics and Data Sharing

We investigated differences in chordal insertion density, area, and mechanical metrics by heart failure, leaflet, insertion site, and their interactions using linear and zero-inflated negative binomial mixed-effects models in R (version 4.6.1, The R Foundation, Vienna, Austria). We used the `lme4, lmerTest, afex, emmeans, glmmTMB, car`, and `DHARMa` packages. The insertion model included valve and scallop as random effects, the area model included valve as a random effect, while the mechanical models included valve and specimen as random effects. Because several variables exhibited right-skewed distributions, we report descriptive statistics as medians and interquartile ranges [25^th^, 75^th^ percentiles] unless otherwise specified. Model transformations were evaluated using residual Q–Q and residual-versus-fitted plots. We defined statistical significance as p*<*0.05 and used the Holm-Bonferroni method to adjust for multiple pairwise comparisons. Statistical model details are provided in Supplement S2. For our mechanical metrics, we included cross-sectional area and peak stress as covariates in the models, which allow us to account for specimen-to-specimen differences in area and peak stress when comparing groups. All raw data and analysis codes are shared openly as described in the Data Availability Statement 8. Lastly, we performed additional statistical investigations between all metrics and the subject data given in Supplementary Table S1. For each outcome, we evaluated age, weight, sex, and valve morphology in separate exploratory models containing one subject-level predictor at a time. Only subject sex was significantly associated with cross-sectional area. We do not report the age, weight, sex, and morphology analyses for any other outcomes.

## 3. Results

### 3.1. Chordal Insertion Density and Rate

We first investigated the number and spatial distribution of chordal insertions across the tricuspid valve leaflets. In total, we marked 5,710 insertions across 454.414 cm^2^ of leaflet tissue, yielding a total pooled insertion density of 12.566 insertions/cm^2^. The average chordal insertion count per valve was 228 ± 61, where raw and summary counts are provided in Supplementary Table S13. For statistical comparisons, we defined the insertion rate as the expected number of chordal insertions per unit leaflet area (insertions/cm^2^), estimated from the conditional negative-binomial component of our zero-inflated model. Insertion rates differed by radial position (p *<* 0.001), circumferential position (p *<* 0.001), leaflet (p *<* 0.001), and there was a significant heart failure-radial position interaction (p *<* 0.001). The main effect of heart failure alone was not significant (p = 0.057). Chordal insertion rates were lowest near the annulus and highest at the free edge (all p *<* 0.001). Insertion rates were lower in the anterior leaflet than in the posterior or septal leaflet (both p *<* 0.001), with no significant difference between the septal and posterior leaflets (p = 0.576). Within radial regions, insertion rates were lower in heart failure than in controls near the annulus (p = 0.025) and in the belly (p = 0.004), but not at the free edge (p = 0.829). Group comparisons by heart failure, leaflet, radial position, and circumferential position are shown in Supplementary Figure S9.

Figure 2 shows the descriptive chordal insertion densities for each 3×3 mesh cell when grouped by the number of scallops per leaflet. Density generally increased from the annulus to the free edge and was concentrated near the scallop center, although some scallops showed higher densities near the boundaries (Figure 2B). Radial-by-circumferential group distributions are shown in Supplementary Figure S10, but these combinations were not statistically compared.

**Figure 2:**
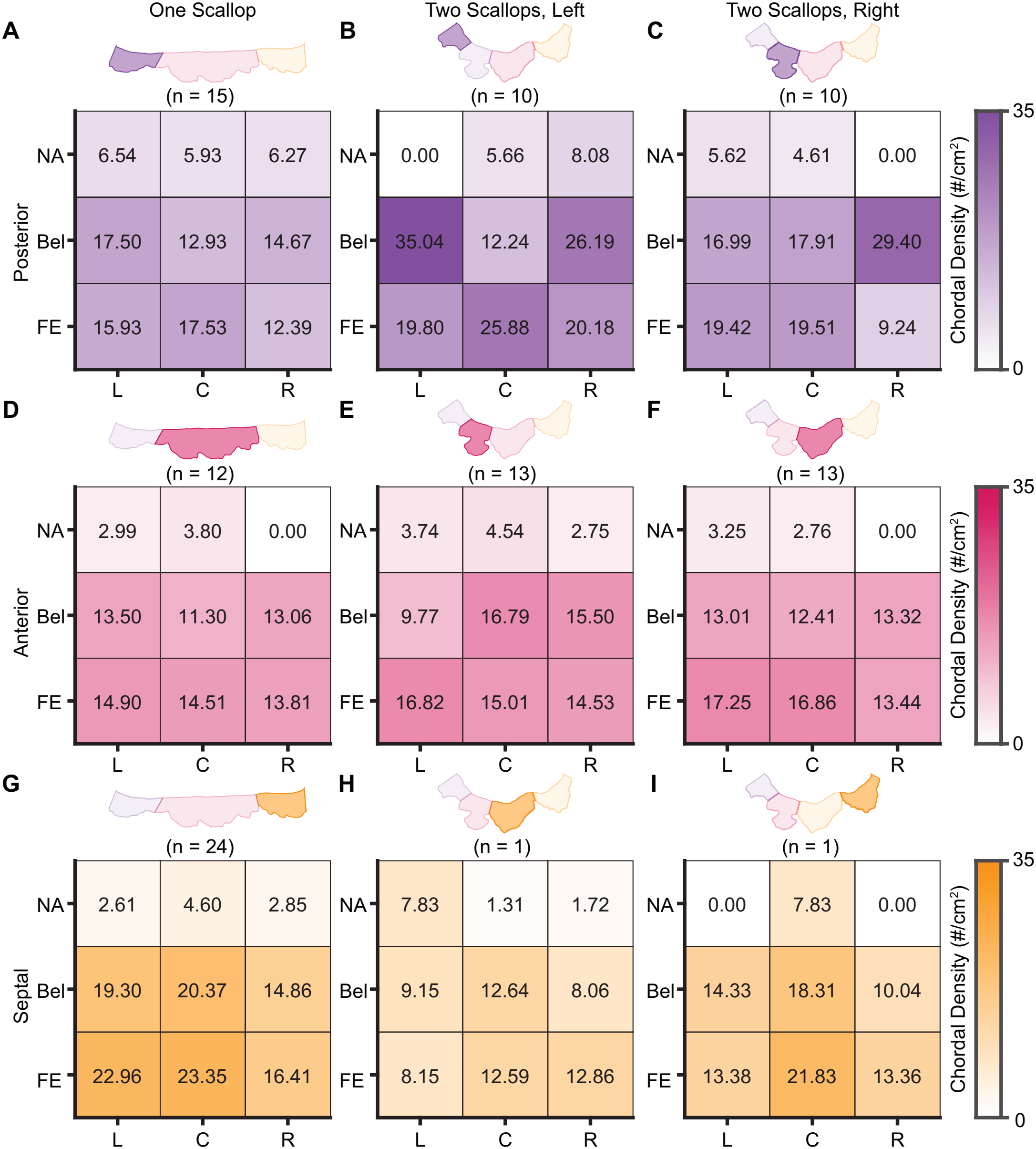
Chordal insertion density differs by leaflet, circumferential, and radial position. Heatmaps show the median chordal insertion density across a 3×3 grid for each scallop. Circumferential regions are left (L), central (C), and right (R), and radial regions are near-annulus (NA), belly (Bel), and free edge (FE). Heatmaps are viewed from the ventricular side. Panels show density for the **(A-C)** posterior, **(D-F)** anterior, and **(G-I)** septal leaflets. For each leaflet, **(A**,**D**,**G)** shows when only one scallop is present, while **(B**,**E**,**H)** is the left scallop and **(C**,**F**,**I)** is the right scallop when two scallops are present. Values are the medians of the cell-level insertion densities, calculated for each scallop as the insertion count divided by the corresponding cell area, where **n** denotes the number of scallops.

### 3.2. Chordal Size

We next investigated the cross-sectional areas of the chordae tendineae. For the 270 excised chordae with insertion site classification, mean cross-sectional area was 0.322 mm^2^, and median was 0.223 [0.132, 0.356] mm^2^. Area depended on leaflet (p *<* 0.001), insertion site (p = 0.003), their interaction (p *<* 0.001), but not on the presence of heart failure (p = 0.941). We grouped the cross-sectional area for each specimen by heart failure, leaflet, and insertion site in Figure 3. Chordae inserting in the anterior leaflet were thicker than those inserting in the posterior and septal leaflets, while chordae inserting in the posterior leaflet were thicker than those inserting in the septal leaflet (all p *<* 0.001, Figure 3B). Further, chordae inserting in the belly region were larger than those inserting in the near-annulus (p = 0.029) or free edge regions (p = 0.004), with no significant difference between near-annulus and free edge (Figure 3C). Lastly, the median cross-sectional area for each leaflet-by-insertion cell is shown in the schematic in Figure 3D. The chordae inserting in the anterior near-annulus and belly regions had the largest median areas, while those inserting in the septal near-annulus and free edge regions had the smallest. Within-group marginal comparisons for leaflet-by-insertion are provided in Supplementary Figure S11. Lastly, we identified an association between sex and chordal cross-sectional area (p = 0.044), with female chordae found to be larger than male (p = 0.046). However, sex was imbalanced across cohorts, with all mechanically tested heart-failure valves being male, so heart-failure status was omitted from the sex model and thus this finding should be interpreted cautiously.

**Figure 3:**
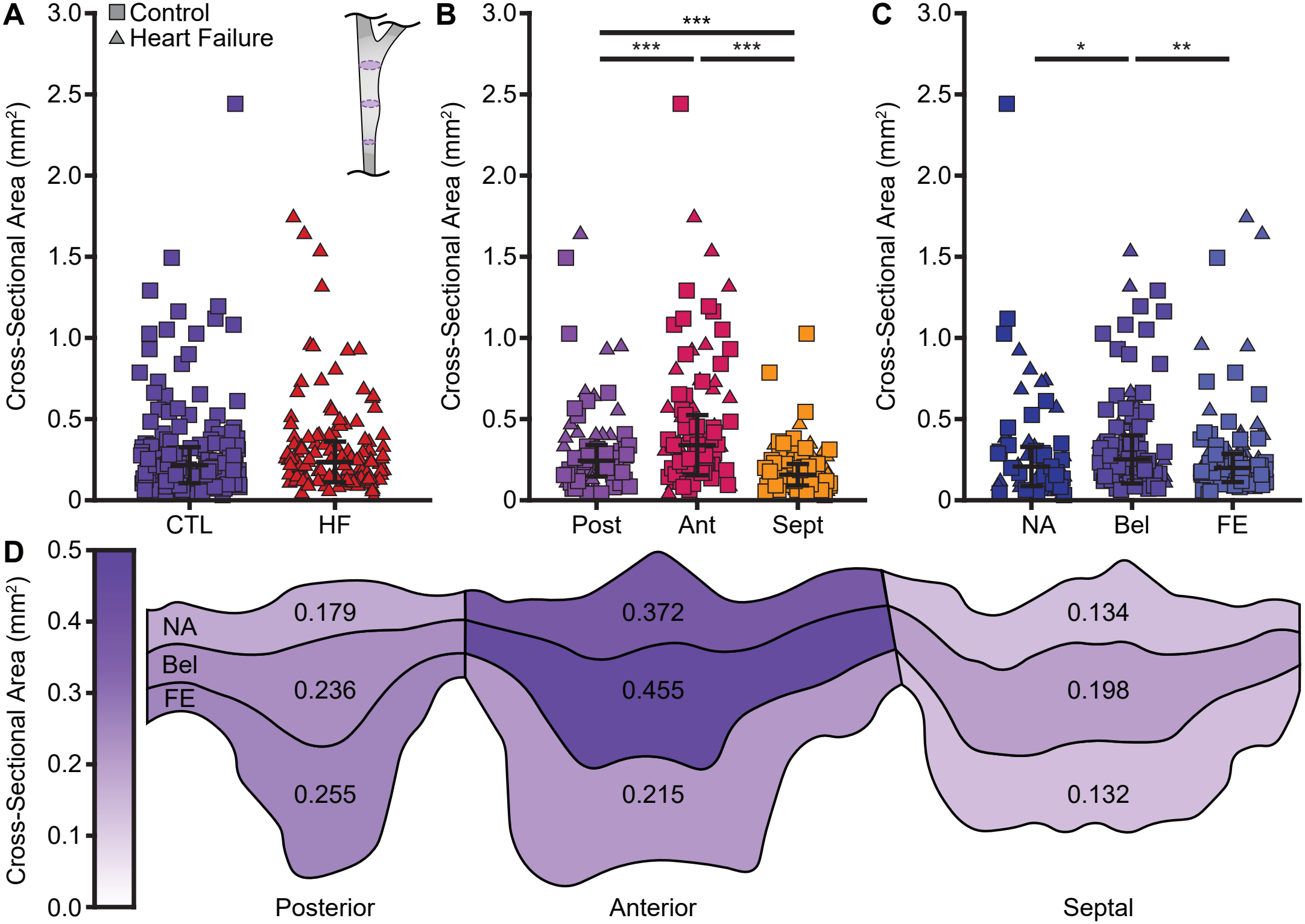
Chordae tendineae cross-sectional area differs by leaflet and insertion site, where the anterior near-annulus and belly are the largest and the septal near-annulus and free edge are the smallest. Cross-sectional area grouped by **(A)** heart failure, **(B)** leaflet, and **(C)** insertion site for 270 specimens, where error bars indicate the median and interquartile range. **(D)** Median cross-sectional areas for the nine leaflet-by-insertion site groups. Areas are computed from the mean diameter measured at three locations assuming a circular cross section (**A** inset). Eleven specimens without insertion site classification are omitted. Asterisks indicate statistical significance (* p*<*0.05, ** p*<*0.01, *** p*<*0.001).

### 3.3. Chordal Mechanics

Finally, we investigated the mechanical properties of the chordae tendineae. The insertion-classified mechanical dataset comprised 689 uniaxial tests on 254 chordal specimens from 11 valves, after excluding 27 specimens with missing insertion site classification or poor testing data. The mean stretch rate was 0.014±0.004 (sec^-1^), with 241 tests to 1 MPa, 232 tests to 2 MPa, and 216 tests to at least 3 MPa. The median stress-stretch curve over the shared stress range of [0, 1] MPa when grouped by heart failure, leaflet, insertion site, and size is shown in Figure 4. The median response varied little by the presence of heart failure, leaflet, or insertion site (Figure 4A-C). Smaller specimens generally exhibited stiffer responses than larger chordae (Figure 4D).

**Figure 4:**
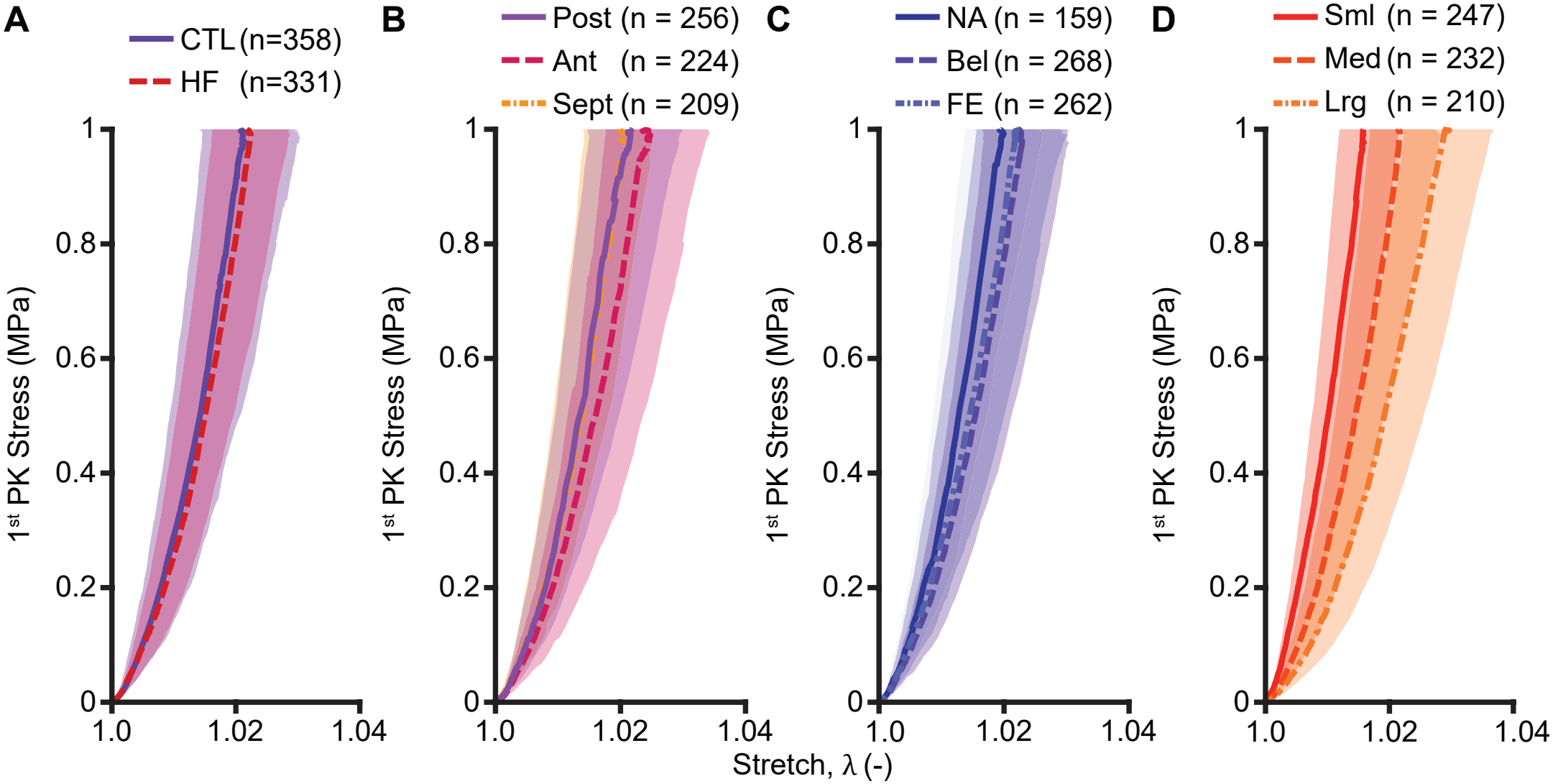
Smaller chordae exhibited stiffer responses than larger chordae. Median stress-stretch responses were grouped by **(A)** heart failure, **(B)** leaflet, **(C)** insertion site, and **(D)** size. Curves show the pointwise median stretch and interquartile range at each stress from 0-1 MPa. Each of the **n** tests contributed over its measured stress range. Consequently, the number of tests contributing to the median decreased at higher stresses.

Median toe stiffness was 18.030 [11.282, 28.244] MPa, median calf stiffness was 80.121 [61.789, 105.844] MPa, and median transition stretch was 1.011 [1.007, 1.016]. The calf stiffness for all tests when grouped by heart failure, leaflet, and insertion site is shown in Figure 5. From the linear mixed-effects model, calf stiffness was associated with cross-sectional area (p *<* 0.001), peak stress (p *<* 0.001), and interactions between heart failure and leaflet (p = 0.006), leaflet and insertion site (p = 0.011), and area-leaflet-insertion site (p = 0.005). The main effects of the presence of heart failure (p = 0.701, Figure 5A), leaflet (p = 0.899, Figure 5B), and insertion site (p = 0.180, Figure 5C) alone were not significant. Pairwise comparisons among condition, leaflet, and insertion site groups showed no significant differences after adjustment for area and peak stress. Within the septal leaflet, however, the near-annulus specimens had higher calf stiffness than belly specimens (p = 0.001).

**Figure 5:**
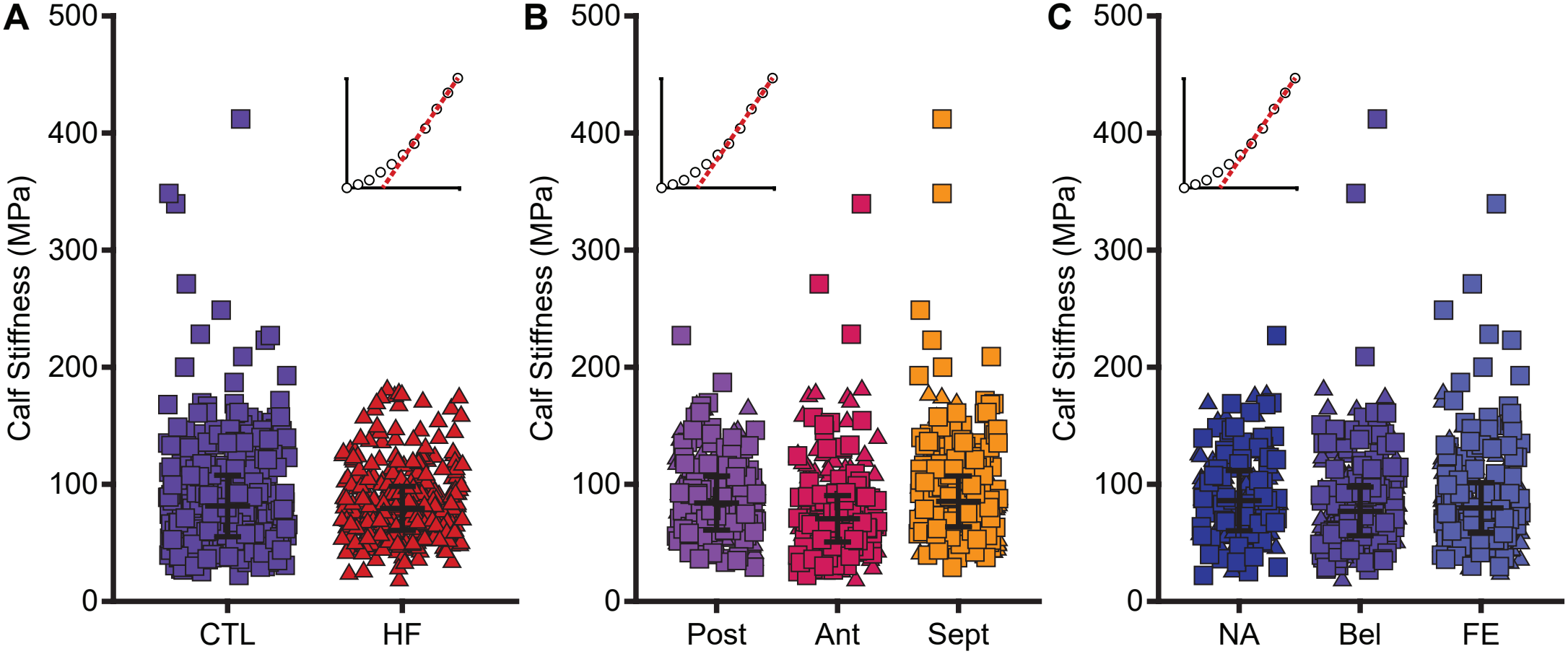
There is no significant difference between chordal calf stiffness when specimens are grouped by **(A)** heart failure, **(B)** leaflet, or **(C)** insertion site after accounting for the cross-sectional area of each specimen. That is, pairwise comparisons among these groups are not significant after adjustment for the cross-sectional area of the specimens. Calf stiffness is defined as the tangent slope near 1 MPa (see insets). Each point represents one of 689 uniaxial tests, where error bars indicate the median and interquartile range. Only tests with a valid insertion site are shown.

Results for toe stiffness and transition stretch, including the effect of peak stress, are provided in Supplement S5. Notably, toe stiffness and transition stretch were also associated with area and peak stress, with no significant main-effect differences among condition, leaflet, or insertion site, although transition stretch was lower in control than in heart failure specimens within the septal leaflet (p = 0.043) and lower in near-annulus than in free edge specimens within the anterior leaflet (p = 0.014). Hysteresis was greater in belly than free edge specimens (p = 0.002), but was not significantly associated with heart failure, leaflet, or cross-sectional area. In summary, toe and calf stiffness decreased with increasing cross-sectional area. After adjustment for area and peak test stress, we detected no overall differences in stiffness by heart failure status, leaflet, or insertion site, although some leaflet-specific contrasts remained.

The mechanical metrics of toe stiffness, calf stiffness, and transition stretch were each associated with specimen cross-sectional area (all p *<* 0.001). Figure 6 shows adjusted relationships estimated from the full mixed-effects models, plotted on the original scales. Area, toe stiffness, and calf stiffness were log-transformed for analysis. Toe and calf stiffness decreased with increasing area. The average marginal log-log slopes were *β* = −0.566[−0.658, −0.474]%*/*% and *β* = −0.351[−0.411, −0.292]%*/*%, for toe and calf stiffness, respectively, averaged across leaflets and insertion sites with the 95% confidence interval values given in brackets. These slopes correspond to approximately 5.3% and 3.3% lower stiffness for a 10% increase in area. On the other hand, transition stretch increased with area (*β* = 0.004[0.003, 0.005]Δ*λ* per unit ln(*A*)). In addition, we found a significant interaction between area, leaflet, and insertion site (p = 0.005) when predicting calf stiffness. We found that only the septal near-annulus specimens had higher estimated stiffness than septal belly specimens at the 75th percentile of area (p = 0.025), where no other comparisons were significant. Calf stiffness predictions for area-leaflet-insertion are shown in Supplementary Figure S13. In summary, chordal stiffness is inversely correlated with the cross-sectional area of the specimen.

**Figure 6:**
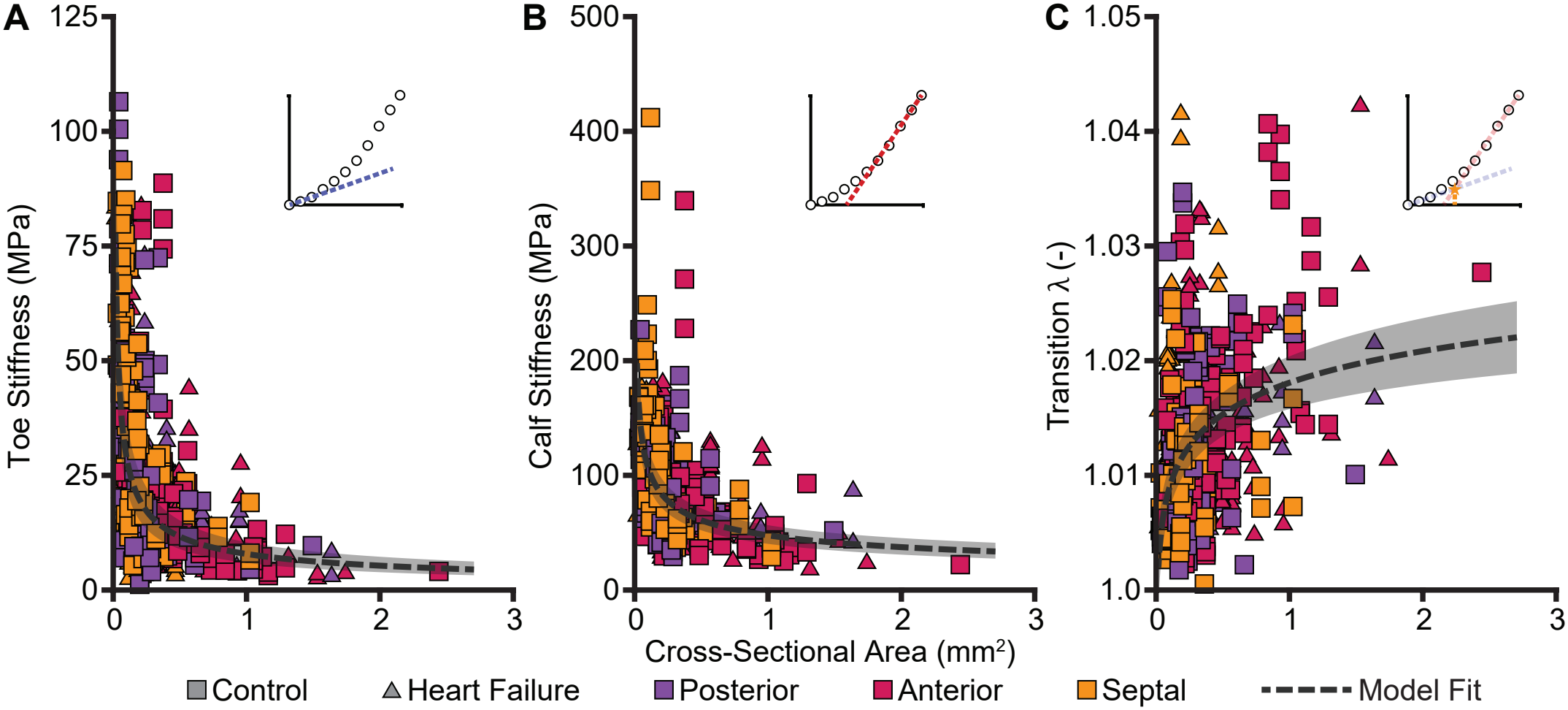
Chordal mechanical behavior is associated with cross-sectional area. **(A)** Toe stiffness, **(B)** calf stiffness, and **(C)** transition stretch are associated with the cross-sectional area for 689 uniaxial tests. Toe and calf stiffness are defined as the tangent slopes near 0 and 1 MPa, respectively. The transition stretch is defined as the intersection of the linear approximations of the toe and calf stiffness, see insets for each. Dashed lines show linear mixed-effects model estimates between the metric and cross-sectional area, with shading showing the 95% confidence intervals.

### 3.4. Chordal Hyperelastic Modeling

We fit ten constitutive models to the experimental stress-stretch data using nonlinear least-squares regression. For each model, we estimated a separate parameter vector for each specimen by fitting its available loading curves jointly. We summarized the specimen-level fits by calculating pooled RMSE and NMSE from the unweighted stress residuals across all specimens, loading curves, and measurement points (Supplementary Eq. S22). The Yeoh-2 and Ogden-1 models fit the data best, with pooled RMSEs of 0.153 and 0.156 MPa, respectively (Supplementary Table S5). Additional terms in the Ogden-2, Ogden-3, and Yeoh-3 models provided little improvement in fit quality. We selected the Ogden-1 model for its fit quality, availability in finite element codes, and to align with prior work on chordae tendineae [20, 19, 13].

In a separate analysis, we used nonlinear least-squares regression to fit the Ogden-1 model to the overall median stress-stretch curve to estimate 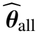 and independently to each group median curve to estimate 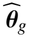. Groups were defined by heart failure, leaflet, insertion site, size, and leaflet-by-insertion site. Supplementary Table S6 reports the resulting Ogden-1 parameters.

We also fit a single hierarchical Bayesian Ogden-1 model to 23,285 stress-stretch observations from 265 chordal specimens. The model estimated specimen-specific material parameters, population-level associations with cross-sectional area, and variation among valves and among specimens within valves. Because cross-sectional area was the only covariate included in the model, the fit included the 11 specimens from the one control valve that lacked insertion-site labels.

To obtain representative population-level Ogden-1 parameters, we set the valve and specimen random effects to zero and evaluated the fitted model at the median cross-sectional area of each leaflet-insertion site group as given in Table 2. These group medians differed slightly from those in Figure 3 because they were calculated only from specimens with usable mechanical data and insertion-site labels. Differences in predicted parameters among groups reflect the different cross-sectional areas at which the same Bayesian model was evaluated. The septal near-annulus and free-edge groups had the highest representative *µ*_1_ and *α*_1_ values, while the anterior near-annulus and belly groups had the lowest. Evaluating the same model at the overall median area (0.224 mm^2^) gave a representative pair of *µ*_1_ = 10.367 MPa, *α*_1_ = 51.569. All reported Ogden-1 parameter sets from least-squares fitting and Bayesian calibration satisfied the Drucker stability criterion and were well-conditioned over the tested stretch range. Additionally, the sensitivity of representative Bayesian parameter estiamtes to sampling settings is reported in Supplement S4.3.

**Table 2:**
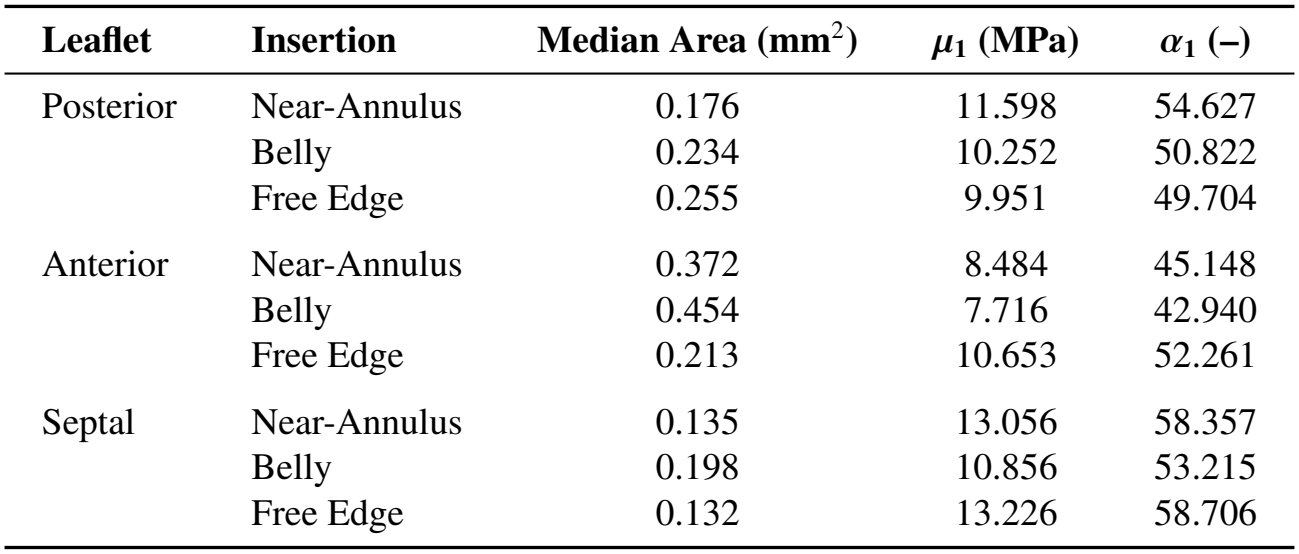
Median cross-sectional areas and representative Ogden-1 parameter pairs (*µ*_1_, *α*_1_) for each leaflet-insertion-site group. At each group’s median area, we evaluated the fitted hierarchical Bayesian model with valve and specimen random effects set to zero and selected the joint posterior draw whose predicted stress-stretch curve was closest to the pointwise posterior median curve.

| **Leaflet** | **Insertion** | **Median Area (mm<sup>2</sup>)** | ***μ*<sub>1</sub> (MPa)** | ***α*<sub>1</sub> (–)** |
|---|---|---|---|---|
| Posterior | Near-Annulus | 0.176 | 11.598 | 54.627 |
|  | Belly | 0.234 | 10.252 | 50.822 |
|  | Free Edge | 0.255 | 9.951 | 49.704 |
| Anterior | Near-Annulus | 0.372 | 8.484 | 45.148 |
|  | Belly | 0.454 | 7.716 | 42.940 |
|  | Free Edge | 0.213 | 10.653 | 52.261 |
| Septal | Near-Annulus | 0.135 | 13.056 | 58.357 |
|  | Belly | 0.198 | 10.856 | 53.215 |
|  | Free Edge | 0.132 | 13.226 | 58.706 |

Figure 7 compares the representative predicted curves with the experimental data for each leaflet-by-insertion group using the parameters given in Table 2. Predictions were generally more compliant for anterior than septal groups, reflecting their different median areas. A comparison between the median response predicted by the Bayesian model and that identified via least-squares fitting is shown in Supplement Figure S8. We found that the Bayesian predictions were more compliant than least-squares fits to group median curves. In summary, the Ogden-1 material parameters for a representative chorda inserting into each leaflet-by-insertion cell of a population-median valve are given in Table 2.

**Figure 7:**
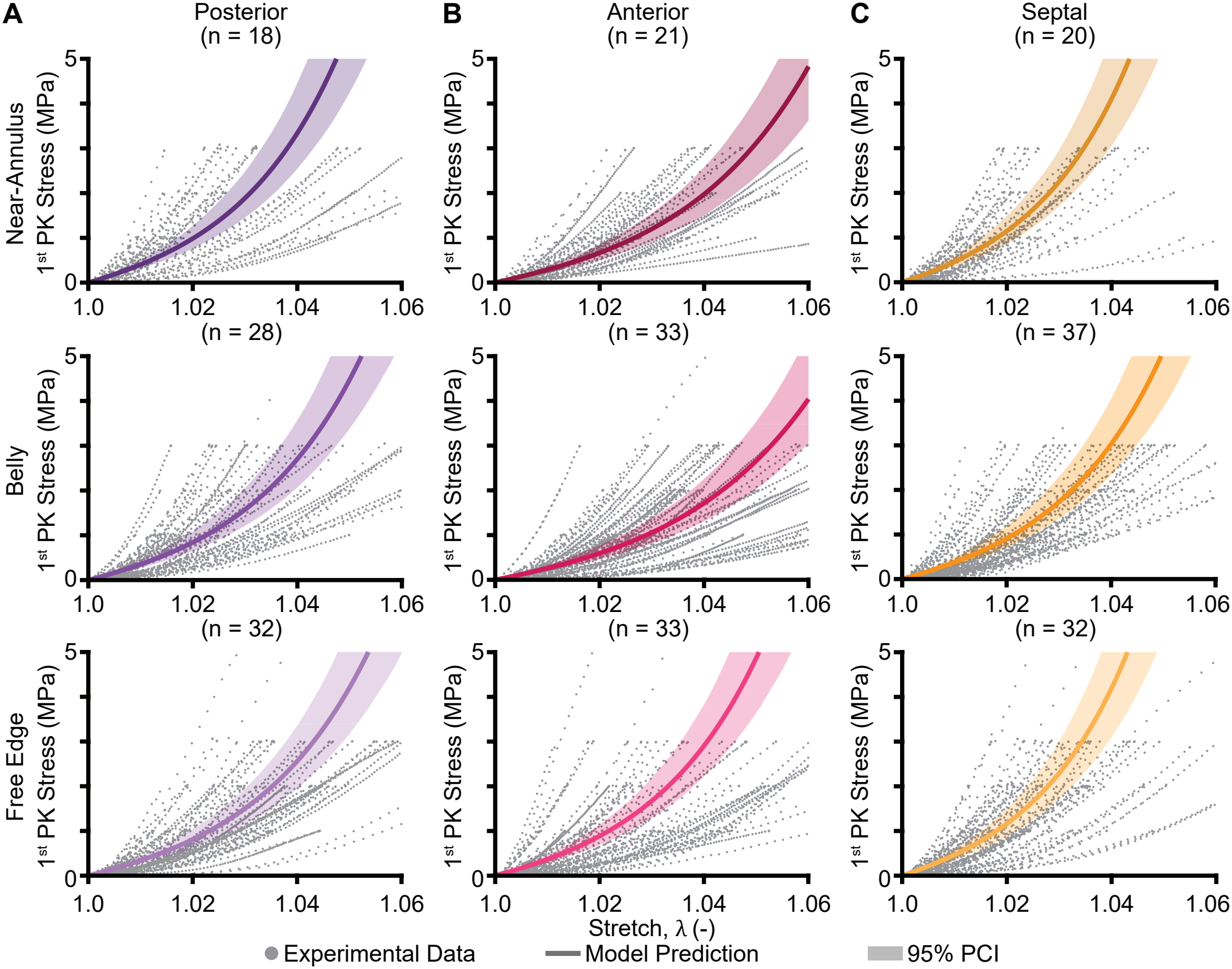
Representative area-conditioned Ogden-1 response of chordae tendineae predicted from the hierarchical Bayesian model. Here, **(A)** is the posterior, **(B)** anterior, and **(C)** septal leaflet, where the rows show the near-annulus, belly, and free edge groups. Curves use the material parameters given in Table 2, and are plotted against experimental data where **n** is the number of specimens. Shading indicates pointwise 95% equal-tailed posterior credible intervals (95% PCI: 2.5^th^–97.5^th^ posterior percentiles) for the population-level stress response at each group’s median cross-sectional area, with valve and specimen random effects set to zero.

## 4. Discussion

Image-based computational models of the TV require information on the chordae tendineae that are often unavailable from clinical images alone. In this study, our primary objective was to characterize human TV chordal insertion distributions, cross-sectional areas, and mechanical properties. Insertion density varied with leaflet and position, while cross-sectional area depended on leaflet, insertion site, and their interaction. Mechanical behavior was strongly associated with chordal size. Smaller chordae exhibited higher material stiffness. Our secondary objective was to test the hypothesis that chordal area and material stiffness were associated with heart failure. Contrary to our hypothesis, neither cross-sectional area nor material stiffness was greater overall in hearts with heart failure than in controls. Together, these findings provide data for assigning chordal geometry and support using a common area-conditioned model to assign material properties in computational models when subject-specific measurements are unavailable.

To our primary objective, we found that chordal insertion density was highest in the belly and free-edge regions, broadly agreeing with prior anatomical descriptions [14, 15, 16]. We identified an average of 228 ± 61 insertions per valve, substantially more than counts reported by Silver et al. (25 [14]) and Skwarek et al. (72.71±13.38 [16]). These discrepancies likely reflect differences in how trunks, branches, and terminal insertions were counted. Our counts represent terminal leaflet attachments rather than independent chordal trunks. The anterior leaflet had the most insertions but a lower insertion density than the posterior or septal leaflet, emphasizing the distinction between total count and density. Regional counts normalized by area can be scaled to leaflet geometry and directly inform computational models.

We also observed that conditional chordal insertion rates were lower in the near-annulus and belly regions of hearts with heart failure when compared to controls, but not overall between the cohorts. This may point to regional leaflet enlargement, which could account for the lower insertion density if attachment counts remained stable [26, 27]. To what extent the heart failure leaflets grew and remodeled was not measured in this study.

Chordal cross-sectional area varied by leaflet, insertion site, and their interaction. In this cohort, the anterior leaflet had the largest chordae and the lowest insertion density. Prior animal studies have also reported size differences among chordal groups [13], although differences in anatomical classifications and measurement methods complicate direct comparisons. The mean cross-sectional area in our cohort (0.322 mm^2^) fell within the range of values reported for human tricuspid chordae, from 0.171 mm^2^ [21] to 0.5 mm^2^ [25] to 1.0 mm^2^ [22].

We found that thinner chordae exhibited higher material stiffness than thicker chordae, consistent with prior human mitral and animal atrioventricular studies [17, 18, 52, 20]. In porcine mitral chordae, Liao and Vesely proposed that smaller collagen fibrils and greater fibril density increased fibril-to-fibril interactions, contributing to the higher material stiffness of smaller chordae. They also reported greater collagen crimp in larger chordae, potentially allowing greater extension before fiber recruitment [52, 53]. These mechanisms may explain our observations, but we did not characterize chordal microstructure in this study. The observed inverse area-stiffness relationship may also promote a more uniform distribution of load among chordae during valve closure. For chordae of similar reference length, higher material stiffness could partly offset the effect of smaller area on resistance to elongation, but this remains to be tested.

Median toe stiffness in our cohort was 18.030 MPa, close to the 16.61 MPa reported by Lim et al.[21] but higher than the 1.73 MPa they later reported for control chordae[22]. Median calf stiffness was 80.121 MPa, within the range of previously reported values for human chordae of 61.8 MPa [25], 85 MPa [22], and 338 MPa [21].

After adjustment for cross-sectional area and peak stress, we found no significant main effects of heart failure status, leaflet, or insertion site on toe or calf stiffness. Because chordal area varied by leaflet and insertion site, differences in size may help explain why anatomical comparisons have yielded different results across studies. Smith et al. found differences in stiffness between leaflets and insertion sites only at small strains [20], whereas Pokutta-Paskaleva et al. reported that both toe and calf stiffness varied by leaflet and insertion site [13]. Some anatomical differences persisted in our cohort after adjustment for area. Within the septal leaflet, near-annulus specimens had higher calf stiffness than belly specimens. Across leaflets, belly specimens also exhibited greater hysteresis than free-edge specimens. Thus, area alone did not explain all variation in measured mechanics.

Among the ten constitutive models we evaluated, the Ogden and Yeoh models provided the best least-squares fits. We selected the one-term Ogden model for the hierarchical Bayesian analysis and estimated its parameters as functions of chordal area while accounting for variation among valves and specimens[50, 51]. Setting valve and specimen effects to zero yields a population-level prediction for a chorda of a specified area. At the median area of the mechanically tested specimens (0.224 mm^2^), the estimated parameters (*µ* = 10.367 MPa, *α* = 51.569) were similar to those we previously reported for a subset of these data (*µ* = 9.03 MPa, *α* = 40.21) [54, 24], but higher than those reported for animal chordae [13, 20, 19].

Regarding our secondary objective, our findings did not support the hypothesis that chordae from hearts with heart failure would be larger and stiffer overall. The association between heart failure and calf stiffness, however, varied by leaflet, suggesting there may be a leaflet-specific impact of health status on chordal mechanics. The recipient hearts came from patients undergoing transplantation for heart failure and did not necessarily have intrinsic TV disease. Our findings therefore do not contradict the reduced stiffness and strength reported in myxomatous tricuspid and mitral chordae [22, 55]. Larger cohorts with defined TV pathologies are needed to better investigate this relationship and determine whether health-specific modeling inputs are needed.

These data are especially useful for image-based tricuspid valve models. The number of terminal attachments in each leaflet region can be estimated by multiplying its surface area by the corresponding insertion density. The chordal cross-sectional areas can then be assigned from the measured distribution for each leaflet and insertion-site group, and these areas can be used to select the corresponding Ogden parameters. This preserves the observed relationship between chordal size and material stiffness. However, these data do not describe the complete chordal architecture. Chordal origins, lengths, branches, fans, and prestress still require additional measurements or assumptions [23, 11]. Future studies should evaluate how these assumptions affect predictive accuracy and mechanics. Lastly, we highlight that all raw testing data are available through the Texas Data Repository [56].

### 4.1. Limitations and Future Work

Our study has limitations. For one, we excised chordae most amenable to mechanical testing. This often favored the largest specimens which may have led to an overestimation of the median cross-sectional area. Furthermore, we assumed the cross-sections of the chordae to be circular and uniform [52, 18]. While cross-sections change along the length of the chordae and are not perfect circles, we aimed to minimize this limitation by measuring the diameter at three locations for each specimen [19]. Because area is both a predictor for mechanical behavior and used to calculate stress, this measurement error could contribute to the observed area-stiffness relationship. Future studies should measure cross-sectional shape and variation more comprehensively to assess the sensitivity of results to this assumption. Moreover, we detected no overall increase in chordal area or stiffness with heart failure. This may have been because the heart failure hearts did not necessarily have TV disease. Additionally, our 281 specimens only came from 12 independent hearts, limiting the population represented. The absence of a female recipient heart among tested valves further limited our ability to draw conclusions about sex and heart failure effects. For the chordal insertion site maps, a single observer segmented all valves; a future investigation should quantify inter-observer variability and its impact on the maps. Finally, we did not quantify chordal length or retain papillary muscle origins. Troxler et al. reported force differences associated with chordal origin [57], motivating further study of how anatomy and *in vivo* loading relate to material behavior. Future work should preserve papillary muscle connections and characterize microstructure to better explain the size-dependent response.

## 5. Conclusion

In this study, we quantified how many chordae insert into a leaflet, their size, and their mechanical properties. We found that insertion density and cross-sectional area varied by anatomical location. Contrary to our hypothesis, we found no overall increase in chordal area or material stiffness associated with heart failure. Smaller specimens exhibited higher material stiffness than large specimens. We used a hierarchical Bayesian model to estimate area-dependent one-term Ogden parameters, providing material inputs that preserve the observed size-stiffness relationship. Together, these data support assigning chordal geometry and material properties in image-based models when subject-specific measurements are unavailable.

## 6. Acknowledgments

This research was supported by funding from the National Heart, Lung, and Blood Institute of the National Institutes of Health through award numbers R01HL165251 and R21HL161832 to M.K.R. and T.A.T., F31HL178280 to C.E.H, and in part by the American Heart Association through award number 25IPA1454136 to W.H.

## 7. Declaration of Competing Interest

M.K.R. has a speaking agreement with Edwards Lifesciences and LumenexBio. All other authors declare that they have no known competing financial interests or personal relationships that could have appeared to influence the work reported in this manuscript.

## 8. Data Availability Statement

All raw mechanical data, strain tracking files, MATLAB processing codes, R codes, and Python codes are available through the Texas Data Repository [56].

## 9. Declaration of generative AI and AI-assisted technologies in the manuscript preparation process

During the preparation of this work the authors used ChatGPT, Claude, and Codex to assist with coding, data processing, drafting, and editing. After using these tools, the authors reviewed and edited the content as needed and take full responsibility for the content of the published article.

## Notes

https://doi.org/10.18738/T8/J9CPSN

### Summary of Updates

Updated the main text of the manuscript. Reworded and added some materials to the Supplement regarding the Bayesian model.

