## Supplement for "Tricuspid Valve Chordae Tendineae: Where are They, How Many are There, and What are Their Properties?"

### S1. Subject-Specific Data

We collected twenty-five human tricuspid valves for this study. Each valve was excised and photos were taken on a calibrated mat, an example of which is shown in Figure S1. Table S1 lists the valvular morphology [1], sex, age, weight, height, and body-surface area (BSA) for each valve. Chordae tendineae from 12 valves were excised and mechanically tested. The subject data for this cohort of valves is given in Table S2.

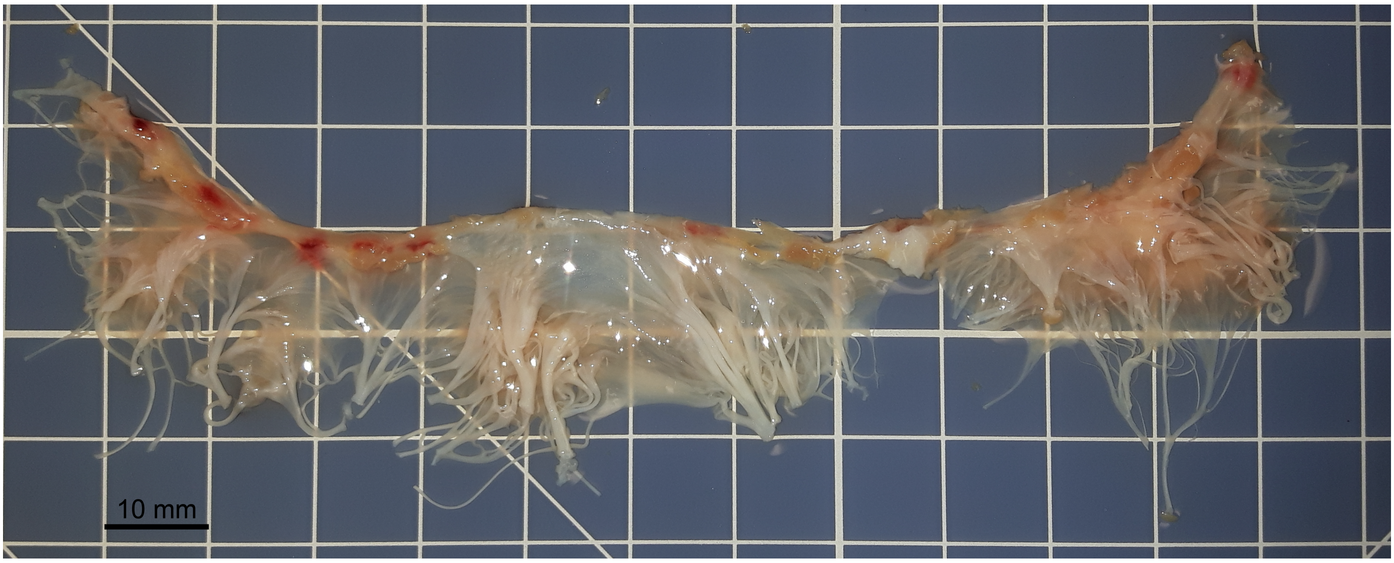


Figure S1: Example photo of an excised tricuspid valve used in this study on a calibrated mat, namely CTL1.

We excised 281 chordae tendineae and recorded their leaflet and radial insertion sites where available. Table S3 provides specimen counts by heart failure, leaflet, and insertion site. Eleven specimens from CTL 3 lacked insertion site information.

*Corresponding Author

Table S1: Subject data for all excised tricuspid valves collected for this study, with average values provided below each cohort of valves. Valve morphology is defined according to Hahn et al. [1].

| Specimen | Morphology | Sex | Age (yrs) | Weight (kg) | Height (cm) | BSA ($m^2$) | TR Grade |
|---|---|---|---|---|---|---|---|
| **Mechanically Tested Valves** | | | | | | | |
| *Control* | | | | | | | |
| CTL1 | IIIB | F | 37 | 69 | 158.0 | 1.74 | ≤ Trace |
| CTL2 | IIIA | F | 59 | 89 | – | – | ≤ Trace |
| CTL3* | IIIA | M | 41 | 108 | 176.3 | 2.3 | ≤ Trace |
| CTL4* | IV | F | 58 | 48 | 151.2 | 1.42 | ≤ Trace |
| CTL5* | IIIA | F | 45 | 74 | 150.0 | 1.76 | ≤ Trace |
| CTL6 | IIIA | M | 20 | 64 | – | – | ≤ Trace |
| CTL7 | I | M | 57 | 95 | 170.0 | 2.12 | ≤ Trace |
| CTL8 | IV | F | 56 | 105 | – | – | ≤ Trace |
| *Average* | | | 46.6 | 81.5 | 161.1 | 1.9 | ≤ Trace |
| *Heart Failure* | | | | | | | |
| HF1 | IV | M | 51 | 94.8 | 162.6 | 2.07 | Moderate |
| HF2 | IV | M | 54 | 126.2 | 175.3 | 2.48 | Mild |
| HF3 | I | M | 26 | 75.9 | 182.9 | 1.96 | Moderate |
| HF4 | I | M | 60 | 58.2 | 162.6 | 1.62 | Severe |
| *Average* | | | 47.8 | 88.8 | 170.9 | 2.0 | – |
| *All Mechanically Tested Valves* | | | | | | | |
| *Average* | | | 47 | 83.9 | 165.4 | 1.9 | – |
| **Additional Valves** (insertion mapping only; no mechanical testing) | | | | | | | |
| *Control* | | | | | | | |
| CTL9 | IIIB | F | 64 | – | – | – | – |
| CTL10 | IIIB | M | 69 | – | – | – | – |
| CTL11 | IIIA | F | 63 | – | – | – | – |
| CTL12 | IIIB | M | 41 | – | – | – | – |
| CTL13 | IIIA | M | 50 | – | – | – | – |
| CTL14 | IIIC | M | 66 | – | – | – | – |
| *Heart Failure* | | | | | | | |
| HF5 | I | F | 24 | 58.2 | 160.0 | 1.61 | Severe |
| HF6 | IIIA | M | 47 | 76.0 | 172.2 | 1.91 | Moderate-Severe |
| HF7 | IIIA | F | 52 | – | – | – | Trace |
| HF8 | I | M | 61 | – | – | – | Severe |
| HF9 | IV | M | 27 | – | – | – | Trace |
| HF10 | IIIB | – | – | – | – | – | – |
| HF11 | I | – | – | – | – | – | |

*Valve was tested on the CellScale UniVert. – Indicates the data is unavailable.

Table S2: Summary of subject data for all valves from which chordae tendineae were mechanically tested. Categorical values are shown as the number, **n** (% of group), and numeric values are shown as the mean ± one standard deviation. CTL = control, HF = heart failure.

| | | **Sex** | | **Age** | **Weight** | **Morphology** | | | | | |
|---|---|---|---|---|---|---|---|---|---|---|---|
| **Group** | **n** | Male | Female | (yrs) | (kg) | I | II | IIIA | IIIB | IIIC | IV |
| All | 12 | 7 (58.3) | 5 (41.7) | 47.0 ± 13.5 | 83.9 ± 23.0 | 3 (25.0) | 0 (0) | 4 (33.3) | 1 (8.3) | 0 (0) | 4 (33.3) |
| CTL | 8 | 3 (37.5) | 5 (62.5) | 46.6 ± 13.7 | 81.5 ± 21.2 | 1 (12.5) | 0 (0) | 4 (50.0) | 1 (12.5) | 0 (0) | 2 (25.0) |
| HF | 4 | 4 (100.0) | 0 (0) | 47.8 ± 15.0 | 88.8 ± 29.1 | 2 (50.0) | 0 (0) | 0 (0) | 0 (0) | 0 (0) | 2 (50.0) |

Table S3: Number of chordae excised when grouped by heart failure, leaflet, and insertion site (total $n = 281$). Bold diagonal entries give the total for each category, off-diagonal entries give the number of chordae falling in both categories. Insertion site was unavailable for eleven control specimens from one valve (CTL 3), so insertion-site counts sum to 270.

| | Heart Failure | | Leaflet | | | Insertion Site | | |
|---|---|---|---|---|---|---|---|---|
| | Control | Heart Failure | Posterior | Anterior | Septal | Near-Annulus | Belly | Free Edge |
| **Heart Failure** | | | | | | | | |
| Control | **160** | | 48 | 58 | 54 | 32 | 61 | 56 |
| Heart Failure | | **121** | 37 | 42 | 42 | 29 | 43 | 49 |
| **Leaflet** | | | | | | | | |
| Posterior | | | **85** | | | 19 | 29 | 34 |
| Anterior | | | | **100** | | 21 | 36 | 38 |
| Septal | | | | | **96** | 21 | 39 | 33 |
| **Insertion Site** | | | | | | | | |
| Near-Annulus | | | | | | **61** | | |
| Belly | | | | | | | **104** | |
| Free Edge | | | | | | | | **105** |

### S2. Details on Statistical Models

#### *S2.1. Chordal Insertion Counts and Density*

For the chordal insertion density, each observation comprised an insertion count and the corresponding cell area within a leaflet scallop's 3×3 grid. A traditional linear mixed-effects model would compare the mean of several cell densities which would unequally weight cells with different areas. To this end, for descriptive comparisons across pooled anatomical regions we instead calculated the "pooled" density as the sum of insertion counts divided by the sum of sampled areas, including zero-count cells. For inference, we modeled counts using a zero-inflated negative-binomial mixed-effects model with log-transformed cell area as an offset. All logarithms denote natural logarithms. Pairwise comparisons used estimated marginal insertion rates from the conditional negative-binomial component (`component = "cond"`), standardized to 1 cm$^2$. These comparisons describe conditional insertion rates rather than overall expected densities incorporating the zero-inflation component.

We compared Poisson, negative-binomial, and zero-inflated negative-binomial mixed-effects models using likelihood-ratio tests, AIC, and BIC. The Poisson and negative-binomial models under-predicted zero counts, so for the primary inferential analysis we selected a zero-inflated negative-binomial model with a quadratic mean-variance relationship (`nbinom2`). Both valve and scallop were included as random effects. Subsequently, we hierarchically tested the conditional fixed-effect interactions while holding the zero-inflation model constant. The final conditional model included heart failure (CTL/HF), leaflet (Ant/Post/Sept), radial position (insertion site: NA/BEL/FE), circumferential position (L/C/R), and heart failure-by-radial position interactions as fixed effects. The zero-inflation component included additive radial and circumferential effects. We assessed fixed effects using Type III Wald tests with sum-to-zero contrasts and adjusted pairwise comparisons using Holm-Bonferroni adjustment.

Let $HF$ denote heart failure group, $Leaf$ leaflet, $Ins$ insertion site (radial position), and $Circ$ circumferential position. Let $N_{vcj}$ denote the insertion count in grid cell $j$ of scallop $c$ within valve $v$, and let $a_{vcj}$ denote that region's area in cm$^2$. The valve random intercept is $b_v$, and the scallop random intercept is $b_{vc}$. The final zero-inflated negative-binomial model is,

$$N_{vcj} \sim \text{ZINB}_2\left(\mu_{vcj}, \theta, \pi_{vcj}\right), \tag{S1}$$

$$\log(\mu_{vcj}) = \log(a_{vcj}) + \beta_0 + \beta_{HF} + \beta_{Leaf} + \beta_{Ins} + \beta_{Circ} + \beta_{HF\cdot Ins} + b_v + b_{vc}, \tag{S2}$$

$$\text{logit}(\pi_{vcj}) = \gamma_0 + \gamma_{Ins} + \gamma_{Circ} \tag{S3}$$

Here, $\pi_{vcj}$ is the probability of belonging to the additional-zero component. The negative-binomial component has variance,

$$\text{Var}_{\text{NB}}(N) = \mu + \frac{\mu^2}{\theta} \tag{S4}$$

and the conditional insertion rate is,

$$\rho_{vcj} = \frac{\mu_{vcj}}{a_{vcj}} \tag{S5}$$

The expected density incorporating both model components is $(1-\pi_{vcj})\rho_{vcj}$. Estimated marginal insertion rates from the conditional count were standardized to a cell area of 1 cm$^2$. Cells with identical radial and circumferential positions share the same modeled zero-inflation probability.

*S2.2. Cross-Sectional Area and Mechanical Metrics*

We analyzed chordal cross-sectional area using a linear mixed-effects model. We included heart failure, leaflet, insertion site, and the interaction of leaflet and insertion site as fixed effects, with valve as a random effect. Before analysis, we log-transformed the cross-sectional area. For the linear mixed models, $\varepsilon$ is the residual error and $A$ is the chordal cross-sectional area. All logarithms denote natural logarithms. The final model is,

$$\log(A_{vs}) = \beta_0 + \beta_{HF} + \beta_{Leaf} + \beta_{Ins} + \beta_{Leaf\cdot Ins} + b_v + \varepsilon_{vs} \tag{S6}$$

The mechanical metrics of toe stiffness, calf stiffness, transition stretch, and hysteresis were also analyzed using a linear mixed-effects model with valve and specimen included as random effects. The addition of specimen offsets accounted for the repeated tests within a given chordal specimen. Fixed effects included heart failure, leaflet, insertion site, log-transformed cross-sectional area, and peak stress (1, 2, or 3 MPa). Peak stress was modeled as a three-level categorical factor. Toe and calf stiffness were log-transformed, but transition stretch and hysteresis were analyzed on their original scales. After performing an analysis of ANOVA's, we also included heart failure-by-leaflet and leaflet-by-insertion site interactions. For only the calf stiffness model, we also included a leaflet-by-insertion-by-area three-way interaction.

Because chordal cross-sectional area is a continuous predictor, we evaluated fixed-effect structures for size and the mechanical metrics using likelihood-ratio comparisons of maximum-likelihood fits. Final models were fitted using restricted maximum likelihood. Fixed effects were assessed using Type III tests with sum-to-zero contrasts. We evaluated marginal means and area slopes using `emmeans` and the Holm-Bonferroni method for multiple pairwise comparisons. Exploratory analysis also investigated age, weight, sex, and valve morphology as fixed effects for predicting the mechanical metrics. In a sensitivity analysis, we also included testing platform (UniVert/Instron) as an additional fixed effect and found this did not materially alter the estimated area-stiffness relationship or the statistical conclusions. Testing platform was therefore omitted from the primary models.

Categorical fixed-effect terms denote the contributions associated with each observation's factor levels, including the corresponding interaction contributions. Sum-to-zero contrasts were used. For compactness, observation-level indices on these contributions are omitted. Let $A_{vs}$ denote the specimen area, $T$ denote the peak stress, and let $r$ index the repeated tests within a specimen, $s$, within a valve, $v$, such that $b_{vs}$ denotes the random intercept. Our final models for toe stiffness ($K_{\text{toe}}$), transition stretch ($\lambda_{\text{tr}}$), and hysteresis ($H$) are,

$$\log(K_{\text{toe},vsr}) = \beta_0 + \beta_{HF} + \beta_{Leaf} + \beta_{Ins} + \beta_{HF\cdot Leaf} + \beta_{Leaf\cdot Ins} + \beta_{\log(A)} \log(A)_{vs} + \beta_T + b_v + b_{vs} + \varepsilon_{vsr}, \tag{S7}$$

$$\lambda_{\text{tr},vsr} = \beta_0 + \beta_{HF} + \beta_{Leaf} + \beta_{Ins} + \beta_{HF\cdot Leaf} + \beta_{Leaf\cdot Ins} + \beta_{\log(A)} \log(A)_{vs} + \beta_T + b_v + b_{vs} + \varepsilon_{vsr}, \tag{S8}$$

$$H_{vsr} = \beta_0 + \beta_{HF} + \beta_{Leaf} + \beta_{Ins} + \beta_{HF\cdot Leaf} + \beta_{Leaf\cdot Ins} + \beta_{\log(A)} \log(A)_{vs} + \beta_T + b_v + b_{vs} + \varepsilon_{vsr}, \tag{S9}$$

The coefficients for each model are estimated independently. For calf stiffness, the model is,

$$\log(K_{\text{calf},vsr}) = \beta_0 + \beta_{HF} + \beta_{Leaf} + \beta_{Ins} + \beta_{HF\cdot Leaf} + \beta_{Leaf\cdot Ins} + \beta_T + \left(\beta_{\log(A)} + \beta_{Leaf\cdot\log(A)} + \beta_{Ins\cdot\log(A)} + \beta_{Leaf\cdot Ins\cdot\log(A)}\right) \log(A)_{vs} + b_v + b_{vs} + \varepsilon_{vsr} \tag{S10}$$

Further, we assumed the following distributions and estimated the variance components separately for each outcome. Random effects and residuals were assumed mutually independent.

$$b_v \sim \mathcal{N}(0, \sigma_v^2), \qquad b_{vs} \sim \mathcal{N}(0, \sigma_s^2), \qquad \varepsilon \sim \mathcal{N}(0, \sigma_e^2) \tag{S11}$$

Area slopes and predictions shown in Figure 6 were estimated from the full mixed-effects models and averaged across leaflet and insertion-site levels.

### S3. Details and Additional Results of Constitutive Fitting

#### *S3.1. Hyperelastic Derivations and Modeling*

For each uniaxial test we normalized the measured force by the undeformed cross-sectional area and used images to point track and resolve the local tissue stretch. Therefore, the experimental response was expressed as the 1st Piola-Kirchhoff (PK) stress, $P_{11}$, for a given axial stretch, $\lambda_1$. For a given hyperelastic strain-energy function, $W(\lambda_1, \lambda_2, \lambda_3)$, expressed in terms of stretch $\lambda_i$ in the $i^{th}$ principal direction, the nominal stress in principal direction $i$ is given by Eq. S12 [2].

$$P_{ii} = \frac{\partial W}{\partial \lambda_i} + \frac{p}{\lambda_i} \tag{S12}$$

The scalar Lagrange multiplier $p$ is added to enforce incompressibility. We assign $\lambda_1 = \lambda$ as the axial stretch. For uniaxial deformation, we assumed stress-free lateral boundary conditions, $P_{22} = P_{33} = 0$, and solved for $p$ to yield Eq. S13.

$$p = -\lambda_2 \frac{\partial W}{\partial \lambda_2} \tag{S13}$$

For isotropic uniaxial deformation, the transverse stretches are equal, $\lambda_2 = \lambda_3$. Combined with the incompressibility constraint, $\lambda_1\lambda_2\lambda_3 = 1$, this yields $\lambda_2 = \lambda_3 = \lambda^{-1/2}$. Thus, Eq. S12 simplifies to Eq. S15.

$$P_{11} = \frac{\partial W}{\partial \lambda_1} - \lambda_2 \lambda_1^{-1} \frac{\partial W}{\partial \lambda_2} \tag{S14}$$

$$= \frac{\partial W}{\partial \lambda_1} - \lambda^{-3/2} \frac{\partial W}{\partial \lambda_2} \tag{S15}$$

For incompressible uniaxial deformation, the strain-energy density can be expressed as a function of the axial stretch alone, $\overline{W}(\lambda) = W(\lambda, \lambda^{-1/2}, \lambda^{-1/2})$. Applying the chain rule and symmetry between the transverse directions gives,

$$\begin{aligned} \frac{d\overline{W}}{d\lambda} &= \frac{\partial W}{\partial \lambda_1} - \frac{1}{2}\lambda^{-3/2}\left(\frac{\partial W}{\partial \lambda_2} + \frac{\partial W}{\partial \lambda_3}\right) \\ &= \frac{\partial W}{\partial \lambda_1} - \lambda^{-3/2}\frac{\partial W}{\partial \lambda_2} = P_{11}. \end{aligned} \tag{S16}$$

We used this relationship to obtain the uniaxial stress-stretch expressions for the constitutive models listed in Supplementary Table S4. Note that we fit the Ogden models using the parameters $\mu_m$ and $\alpha_m$, where $\mu_m = C_{2m-1}C_{2m}/2$, and $\alpha_m = C_{2m}$ [3, 4]. Thus, the PK stress under incompressible uniaxial deformation was given as,

$$P_{11}(\lambda) = \sum_{m=1}^{N} \frac{2\mu_m}{\alpha_m} \left(\lambda^{\alpha_m - 1} - \lambda^{-\alpha_m/2-1}\right) \tag{S17}$$

where $\mu_m$ has units of MPa and $\alpha_m$ is dimensionless. Furthermore, the one-dimensional linear-elastic model was defined as $P_{11}(\lambda) = E(\lambda - 1)$ and used for comparison. We note that because incompressibility was enforced through the assumed deformation, no volumetric penalty terms were included in the hyperelastic strain-energy functions.

#### *S3.2. Individual Specimen Fits*

For each constitutive model, we aimed to identify the parameter vector $\boldsymbol{\theta}$ bounded by lower and upper bounds $[\boldsymbol{\theta_L}, \boldsymbol{\theta_U}]$ which best minimized the error, $e$, between the experimentally-measured stress, $P^{\text{exp}}$, and the model-computed stress, $P$, where $P \equiv P_{11}$. This optimal constitutive model parameter vector is denoted $\widehat{\boldsymbol{\theta}}$. We used MATLAB's bounded nonlinear least-squares optimizer with 20 multi-start initializations in a custom script to identify $\widehat{\boldsymbol{\theta}}$. First, we fit all specimens individually. Each specimen received one parameter set fit simultaneously to all available loading curves. That is, for specimen $i$, repetition $r$, and measurement point $k$, the stress residual used in the least-squares optimization was defined as,

$$e_{irk}(\boldsymbol{\theta}) = P(\lambda_{irk}; \boldsymbol{\theta}) - P_{irk}^{\text{exp}} \tag{S18}$$

Thus, the fitted specimen-level parameters were

$$\widehat{\boldsymbol{\theta}}_i = \underset{\boldsymbol{\theta}_{\text{L}} \le \boldsymbol{\theta} \le \boldsymbol{\theta}_{\text{U}}}{\arg\min}\, J_i(\boldsymbol{\theta}), \qquad J_i(\boldsymbol{\theta}) = \frac{1}{R_i} \sum_{r=1}^{R_i} \sum_{k=1}^{n_{ir}} w_{irk}^{(\lambda)} e_{irk}^2(\boldsymbol{\theta}) \tag{S19}$$

Table S4: The strain-energy density functions and parameter bounds used for constitutive fitting. All hyperelastic models assume incompressible uniaxial deformation with axial stretch $\lambda$ and transverse stretches $\lambda^{-1/2}$. The axial first Piola-Kirchhoff stress is calculated as $P_{11} = dW/d\lambda$. For compactness, $A(\lambda) = \lambda^2 + 2\lambda^{-1} - 3$ and $B(\lambda) = 2\lambda + \lambda^{-2} - 3$. The linear-elastic model is a one-dimensional Hookean approximation.

| **Model** | **Strain Energy** $W(\lambda)$ | **Parameter Bounds** |
|---|---|---|
| Neo-Hookean | $C_1 A(\lambda)$ | $C_1 \in [10^{-6}, 10^6]$ |
| Fung | $\frac{C_1}{C_2}\left[e^{C_2 A(\lambda)} - 1\right]$ | $C_1 \in [10^{-6}, 10^6]$ <br> $C_2 \in [10^{-9}, 10^4]$ |
| Mooney–Rivlin | $C_1 A(\lambda) + C_2 B(\lambda)$ | $C_1, C_2 \in [0, 10^6]$ |
| Yeoh-2 | $C_1 A(\lambda) + C_2 A(\lambda)^2$ | $C_1 \in [10^{-9}, 10^6]$ <br> $C_2 \in [0, 10^6]$ |
| Ogden-1 | $\sum_{m=1}^{1} \frac{2\mu_m}{\alpha_m^2}\left(\lambda^{\alpha_m} + 2\lambda^{-\alpha_m/2} - 3\right)$ | $\mu_1 \in [10^{-9}, 10^6]$ <br> $\alpha_1 \in [10^{-3}, 10^3]$ |
| Neo-Fung | $C_1 A(\lambda) + \frac{C_2}{2C_3}\left[e^{C_3 A(\lambda)} - 1\right]$ | $C_1, C_2 \in [10^{-9}, 10^6]$ <br> $C_3 \in [10^{-9}, 10^4]$ |
| Yeoh-3 | $C_1 A(\lambda) + C_2 A(\lambda)^2 + C_3 A(\lambda)^3$ | $C_1 \in [10^{-9}, 10^6]$ <br> $C_2 \in [0, 10^6]$ <br> $C_3 \in [0, 10^9]$ |
| Ogden-2 | $\sum_{m=1}^{2} \frac{2\mu_m}{\alpha_m^2}\left(\lambda^{\alpha_m} + 2\lambda^{-\alpha_m/2} - 3\right)$ | $\mu_{1,2} \in [10^{-9}, 10^6]$ <br> $\alpha_{1,2} \in [10^{-3}, 10^3]$ |
| Ogden-3 | $\sum_{m=1}^{3} \frac{2\mu_m}{\alpha_m^2}\left(\lambda^{\alpha_m} + 2\lambda^{-\alpha_m/2} - 3\right)$ | $\mu_{1,2,3} \in [10^{-9}, 10^6]$ <br> $\alpha_{1,2,3} \in [10^{-3}, 10^3]$ |
| Linear Elastic (1D) | $\frac{1}{2}E(\lambda - 1)^2$ | $E \in [10^{-9}, 10^6]$ |

where $R_i$ is the number of repeated loading curves for specimen $i$ and the weights, $w_{irk}^{(\lambda)}$, for each point are chosen such that $\sum_{k=1}^{n_{ir}} w_{irk}^{(\lambda)} = 1$. Here, $n_{ir}$ is the number of measured data points in repetition $r$ of specimen $i$. We chose trapezoidal quadrature weights normalized over the measured stretch interval. Under this construction, every repetition contributed equally regardless of its number of observations or measured stretch range. This prevented curves with more sampled points or spanning larger stretch intervals from disproportionately influencing the fit. For an ordered grid $x_1, \ldots, x_n$, where in this case $x = \lambda$, the unnormalized trapezoidal weights were,

$$\widetilde{w}_k^{(x)} = \begin{cases} \dfrac{x_2 - x_1}{2}, & k = 1, \\ \dfrac{x_{k+1} - x_{k-1}}{2}, & 1 < k < n, \\ \dfrac{x_n - x_{n-1}}{2}, & k = n, \end{cases} \qquad w_k^{(x)} = \frac{\widetilde{w}_k^{(x)}}{\sum_{\ell=1}^{n} \widetilde{w}_\ell^{(x)}} \tag{S20}$$

This weighted residual sum thus approximates the mean squared stress error over the measured stretch, such that,

$$\sum_{k=1}^{n_{ir}} w_{irk}^{(\lambda)} e_{irk}^2 \approx \frac{1}{\lambda_{ir}^{\max} - \lambda_{ir}^{\min}} \int_{\lambda_{ir}^{\min}}^{\lambda_{ir}^{\max}} e_{ir}^2(\lambda)\, \mathrm{d}\lambda \tag{S21}$$

The trapezoidal weights were normalized over the stretch and the stretch values themselves were not rescaled or normalized. Therefore, as defined above in Eqs. (S19)–(S21), each specimen-level parameter vector minimized the mean squared stress error averaged first over the stretch interval of each repetition and then across repetitions. We quantified fit quality using the root-mean-square error (RMSE) and normalized mean-square-error goodness of fit (NMSE) for each individual specimen fit. We further aggregated all error residuals to compute an across-specimen pooled RMSE and NMSE given in Eq. S22, which gives a single summary value of the constitutive model's accuracy and ability to fit the data. A perfect fit has RMSE = 0 and NMSE = 1, and $\overline{P}_i$ is the mean experimental stress across all repetitions and points for specimen $i$.

$$\mathrm{RMSE}_{\mathrm{pooled}} = \sqrt{\frac{\sum_i \sum_r \sum_k e_{irk}^2}{\sum_i \sum_r n_{ir}}} \qquad \mathrm{NMSE}_{\mathrm{pooled}} = 1 - \frac{\sum_i \sum_r \sum_k e_{irk}^2}{\sum_i \sum_r \sum_k \left(P_{irk}^{\exp} - \overline{P}_i\right)^2} \tag{S22}$$

The pooled and average RMSE and NMSE for all ten constitutive models are summarized in Table S5. The elastic, Neo-Hookean, and Mooney-Rivlin models produced the largest errors. The other models produced similar fit qualities to each other. Interestingly, the higher term Ogden and Yeoh models did not noticeably improve fit quality. We note that while the error metrics appear equal, they are rounded to three decimals and the full values differ slightly. Additionally, the parameter bounds allow the higher order terms to equal zero (Yeoh-3: $C_3 = 0$) or approximately equal zero (Ogden: $\mu_2, \mu_3 = 1 \times 10^{-9}$). One optimizer start was initialized at this bound. This suggests that additional terms in the higher order models are unnecessary and converge to zero because they cannot meaningfully reduce the residuals. With this in mind, the Yeoh-2 model had the best fit with a pooled RMSE of 0.153 MPa. The Ogden-1 model was nearly as good, with a pooled RMSE of 0.156 MPa.

Table S5: Root mean squared error (RMSE) and normalized mean squared error (NMSE) for all constitutive models fit to all specimens individually. Pooled errors combine stress residuals across all specimens, repeated tests, and testing points. Average (Avg.) errors are the mean of the 265 specimen-level errors.

| **Model** | **Pooled RMSE (MPa)** | **Pooled NMSE (-)** | **Avg. RMSE (MPa)** | **Avg. NMSE (-)** |
|---|---|---|---|---|
| Neo-Hookean | 0.355 | 0.865 | 0.319 | 0.851 |
| Fung | 0.171 | 0.969 | 0.137 | 0.968 |
| Mooney-Rivlin | 0.355 | 0.865 | 0.319 | 0.851 |
| Yeoh-2 | 0.153 | 0.975 | 0.115 | 0.974 |
| Ogden-1 | 0.156 | 0.974 | 0.120 | 0.973 |
| Neo-Fung | 0.171 | 0.969 | 0.137 | 0.968 |
| Yeoh-3 | 0.153 | 0.975 | 0.115 | 0.974 |
| Ogden-2 | 0.156 | 0.974 | 0.120 | 0.973 |
| Ogden-3 | 0.156 | 0.974 | 0.120 | 0.973 |
| Elastic | 0.343 | 0.874 | 0.308 | 0.860 |

Figure S2 shows the fits and RMSE against the experimental stress-stretch data for one representative specimen for all ten models. The Ogden, Yeoh, and Fung models fit the data well. The Neo-Hookean, Mooney-Rivlin, and linear elastic models were unable to reproduce the non-linear stress-stretch behavior. The Yeoh-2 model had a slightly lower RMSE and higher NMSE than the Ogden-1. We selected the Ogden-1 model for subsequent modeling as described in the main text. Specimen-level parameters are available in the raw data files, see Data Availability Statement.

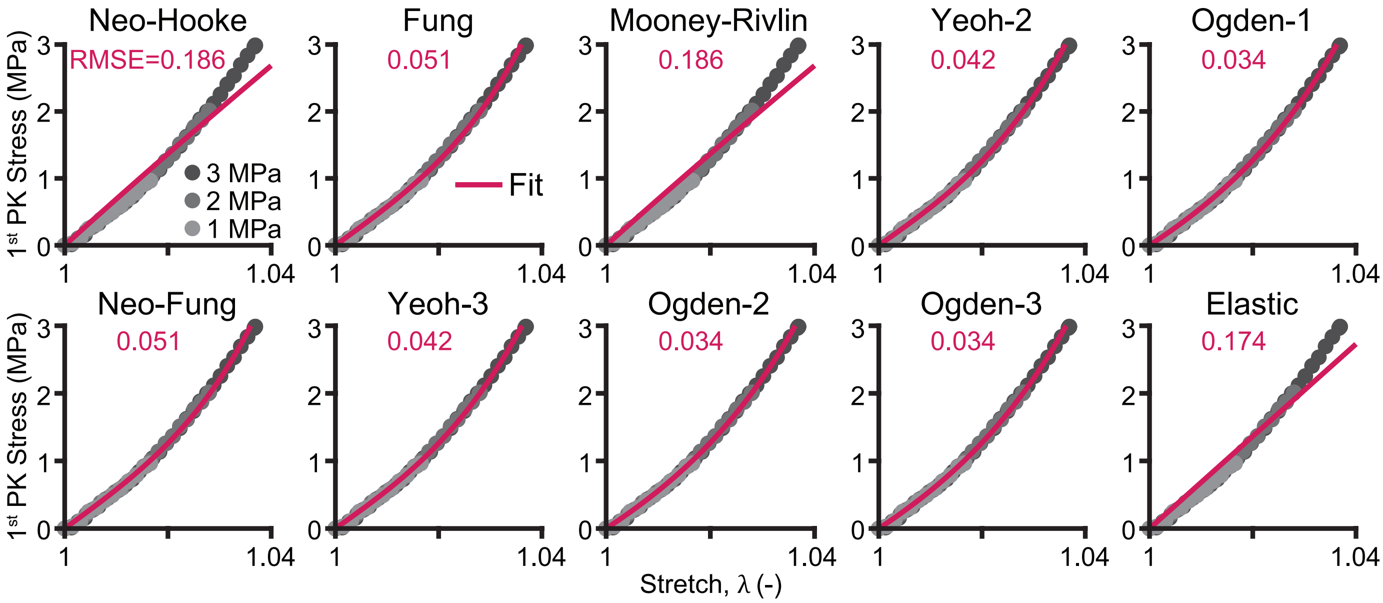


Figure S2: Ten constitutive fits shown against the experimentally measured stress-stretch data with three repeated tests (1, 2, and 3 MPa peak stress) for one specimen selected from the dataset. Root-mean-squared errors (RMSE) in MPa are also given for each fit. The Ogden model produces the best fit with the lowest RMSE for this specimen. The Yeoh and Fung models also produce good fits, while the Neo-Hookean and elastic models are unable to capture the classic non-linear 'J'-shaped stress-stretch curve of the chordae tendineae. The parameters for each model are identified using least-squares fitting to each specimen's test data individually.

*S3.3. Grouped Fits*

We fit representative responses for all specimens combined (all) and for groups defined by heart failure, leaflet, insertion site, size, and leaflet-by-insertion site. For each grouping, we only included specimens which reached at least 1 MPa. First, we added the stress-free reference point $(\lambda, P) = (1, 0)$ and linearly interpolated each stress-stretch curve onto $M = 101$ uniformly-spaced stress points between 0 and 1 MPa. Next, we combined repeated tests in a specimen by taking the pointwise median stretch value across the interpolated stress levels. That is, the representative stretch for specimen $i$ was first calculated from its $R_i^*$ repetitions reaching 1 MPa,

$$\widetilde{\lambda}_i(P_j) = \underset{r \in R_i^*}{\text{median}} \left\{ \lambda_{ir}(P_j) \right\} \tag{S23}$$

Then, we created the median response for the group, $g$, by taking the pointwise median of the specimen-median stretch values across the interpolated stress levels,

$$\widetilde{\lambda}_g(P_j) = \underset{i \in g}{\text{median}} \left\{ \widetilde{\lambda}_i(P_j) \right\} \tag{S24}$$

Thus, $\widetilde{\lambda}_g$ provides one representative stress-stretch curve for each group. We then fit a single parameter set to this median response curve for each model using the same bounded nonlinear least-squares regression (`lsqnonlin`) with 20 multi-start initializations to identify the optimum parameter vector, $\widehat{\boldsymbol{\theta}}_g$, which minimized the normalized, stress-interval weighted objective, $J_g(\boldsymbol{\theta})$. Note that $\widehat{\boldsymbol{\theta}}_{\text{all}}$ includes all specimens in one group. We chose to weight across the stress interval to prevent differences in the original sampling density, number of repeated tests, or stretch range from disproportionately influencing the representative fit. Therefore, the objective is given as,

$$\widehat{\boldsymbol{\theta}}_g = \underset{\boldsymbol{\theta}_{\text{L}} \leq \boldsymbol{\theta} \leq \boldsymbol{\theta}_{\text{U}}}{\arg\min}\, J_g(\boldsymbol{\theta}), \qquad \text{where} \tag{S25}$$

$$J_g(\boldsymbol{\theta}) = \sum_{j=1}^{M} w_j^{(P)} \left[ P\left(\widetilde{\lambda}_g(P_j); \boldsymbol{\theta}\right) - P_j \right]^2, \qquad \sum_{j=1}^{M} w_j^{(P)} = 1 \tag{S26}$$

Here, $w_j^{(P)}$ denotes the stress-normalized trapezoidal weight obtained from Eq. (S20) using $x = P \in [0, 1]$ MPa. Consequently,

$$J_g(\boldsymbol{\theta}) \approx \frac{1}{P_{\text{max}}} \int_0^{P_{\text{max}}} \left[ P\left(\widetilde{\lambda}_g(P); \boldsymbol{\theta}\right) - P \right]^2 \, \text{d}P \tag{S27}$$

Because the stress grid was uniform, the interior points received equal weight, while the two endpoints received one-half of the interior-point weight. For the representative group response, the unweighted goodness-of-fit metrics were calculated after parameter estimation against the $M = 101$-point median group response as,

$$\text{RMSE} = \sqrt{\frac{1}{M} \sum_{j=1}^{M} e_j^2} \qquad \text{NMSE} = 1 - \frac{\sum_{j=1}^{M} e_j^2}{\sum_{j=1}^{M} \left(P_j - \overline{P}\right)^2} \tag{S28}$$

where

$$e_j = P\left(\widetilde{\lambda}_g(P_j); \widehat{\boldsymbol{\theta}}_g\right) - P_j \tag{S29}$$

and $\overline{P}$ is the mean stress. Accordingly, RMSE = 0 and NMSE = 1 indicate a perfect fit. The fitted Ogden parameters for the median responses for each group are provided in Supplement Table S6. The largest difference in Ogden parameters is seen when grouped by size. Differences are also observed when grouped by leaflet and insertion site, e.g. septal free edge versus anterior belly. Parameters are not substantially different when grouped by heart failure, leaflet, and insertion site when compared to the fit of all specimens combined.

Table S6: Ogden-1 parameters estimated by least-squares fitting to median stress-stretch curves. The "All" row reports the fit across all specimens ($\widehat{\boldsymbol{\theta}}_{\text{all}}$); each remaining row reports a separate fit for group $g$ ($\widehat{\boldsymbol{\theta}}_g$). Here, **n** is the number of specimens contributing to the median curve. RMSE and NMSE were calculated by comparing the fitted response with that median curve at 101 points.

| **Group** | **n =** | $\mu_1$ **(MPa)** | $\alpha_1$ **(-)** | $C_1$ **(MPa)** | **RMSE (MPa)** | **NMSE (-)** |
|---|---|---|---|---|---|---|
| All | 265 | 7.053 | 92.162 | 0.153 | 0.0113 | 0.998 |
| Control | 148 | 7.316 | 93.897 | 0.156 | 0.0114 | 0.998 |
| Heart Failure | 117 | 6.947 | 89.327 | 0.156 | 0.0139 | 0.998 |
| Anterior | 92 | 6.474 | 76.693 | 0.169 | 0.0148 | 0.997 |
| Posterior | 81 | 8.125 | 77.894 | 0.209 | 0.0219 | 0.994 |
| Septal | 92 | 8.350 | 91.138 | 0.183 | 0.0203 | 0.995 |
| Near-Annulus | 59 | 7.701 | 99.783 | 0.154 | 0.0161 | 0.997 |
| Belly | 98 | 6.692 | 89.586 | 0.149 | 0.0115 | 0.998 |
| Free Edge | 97 | 7.023 | 90.243 | 0.156 | 0.0110 | 0.999 |
| Small | 88 | 12.374 | 90.241 | 0.274 | 0.0132 | 0.998 |
| Medium | 88 | 7.004 | 93.568 | 0.150 | 0.0128 | 0.998 |
| Large | 89 | 5.681 | 66.891 | 0.170 | 0.0153 | 0.997 |
| Posterior: Near-Annulus | 18 | 9.421 | 120.541 | 0.156 | 0.0135 | 0.998 |
| Posterior: Belly | 28 | 8.230 | 105.247 | 0.156 | 0.0156 | 0.997 |
| Posterior: Free Edge | 32 | 5.332 | 98.890 | 0.108 | 0.0107 | 0.999 |
| Anterior: Near-Annulus | 21 | 6.108 | 96.423 | 0.127 | 0.0133 | 0.998 |
| Anterior: Belly | 33 | 5.684 | 82.912 | 0.137 | 0.0171 | 0.997 |
| Anterior: Free Edge | 33 | 6.234 | 73.399 | 0.170 | 0.0107 | 0.999 |
| Septal: Near-Annulus | 20 | 7.619 | 110.303 | 0.138 | 0.0119 | 0.998 |
| Septal: Belly | 37 | 6.706 | 96.606 | 0.139 | 0.0159 | 0.997 |
| Septal: Free Edge | 32 | 9.113 | 107.939 | 0.169 | 0.0098 | 0.999 |

### S4. Details and Additional Results of Hierarchical Bayesian Model

We fit an area-dependent hierarchical Bayesian model to 23,285 stress-stretch observations from 265 chordal specimens nested within 12 valves. The model estimated the one-term Ogden parameters, $\mu_1$ and $\alpha_1$, as functions of specimen cross-sectional area, with random effects for valves and for specimens within valves. The undeformed cross-sectional area (Area, or $A$) was measured for each chordal specimen and represented as a continuous variable. The distribution of chordae cross-sectional area was right-skewed and differed across leaflet-by-insertion sites. We log-transformed the cross-sectional area before analysis. Based on our statistical analysis and results, we included log-transformed cross-sectional area as the sole covariate in the hierarchical model. Leaflet, insertion site, and heart failure status were not included as predictors.

Following Section 2.5, we assumed incompressible uniaxial extension, with axial stretch $\lambda$ and transverse stretches $\lambda^{-1/2}$. Again, we selected the Ogden-1 model for the hierarchical analysis. For specimen $i$, the first Piola-Kirchhoff axial stress $P_{11} \equiv P$ is,

$$P_i(\lambda) = f(\lambda; \boldsymbol{\theta}_i) = \frac{2\mu_{1i}}{\alpha_{1i}} \left( \lambda^{\alpha_{1i}-1} - \lambda^{-\alpha_{1i}/2-1} \right), \qquad \boldsymbol{\theta}_i = (\mu_{1i}, \alpha_{1i}). \tag{S30}$$

Let $V[i]$ denote the valve containing specimen $i$, and let $k = 1, 2$ index the material parameter, with $\theta_{1i} = \mu_{1i}$ and $\theta_{2i} = \alpha_{1i}$. All logarithms denote natural logarithms. We standardized log cross-sectional area as

$$x_i = \frac{\log A_i - \overline{\log A}}{s_{\log A}}, \qquad \overline{\log A} = -1.4390063, \qquad s_{\log A} = 0.8149903, \tag{S31}$$

where area was expressed in $\text{mm}^2$ before transformation. These training-data standardization constants were also used for predictions. The material parameters followed the non-centered hierarchy,

$$\log \theta_{ki} = \log m_k + \beta_{k,\log A} x_i + \sigma_{k,V} z_{k,V[i]} + \sigma_{k,S} z_{k,i}, \qquad z_{k,V}, z_{k,i} \sim \mathcal{N}(0, 1). \tag{S32}$$

Here, $m_k$ is the population median material parameter at $x_i = 0$, and $\beta_{k,\log A}$ is the change in log material parameter per standard-deviation increase in log area. Thus, $\exp(\beta_{k,\log A})$ gives the corresponding multiplicative change in the material parameter. The term $\sigma_{k,V} z_{k,V[i]}$ is shared by specimens from the same valve, while $\sigma_{k,S} z_{k,i}$ describes the additional departure of specimen $i$ from its valve-level expectation. The scales $\sigma_{k,V}$ and $\sigma_{k,S}$ are the between-valve and within-valve between-specimen standard deviations of each log material parameter. Smaller scales imply stronger pooling. Exponentiating the linear predictor ensures positive material parameters.

For stress observation $j$ from specimen $i$, we used the Gaussian observation model

$$y_{ij} \mid \boldsymbol{\theta}_i, \sigma_{m,V[i]} \sim \mathcal{N}\Big(f(\lambda_{ij}; \boldsymbol{\theta}_i), \sigma^2_{m,V[i]}\Big). \tag{S33}$$

All observations from a specimen shared its constitutive parameters. Each of the twelve valves had its own residual stress standard deviation, $\sigma_{m,V}$, shared across its specimens. This scale captures experimental noise and mismatch between the constitutive response and observations, and is distinct from the between-valve material-parameter scale $\sigma_{k,V}$. Residual stress errors were assumed conditionally independent, and each observation contributed equally to the likelihood. The effects of residual correlation within loading curves and differences in sampling density on parameter estimates and posterior uncertainty were not evaluated.

Table S7 gives the priors for the final model. Normal distributions are parameterized as $\mathcal{N}(\text{mean}, \text{SD}^2)$, and exponential distributions are parameterized by their rate. The hierarchy-scale priors therefore have mean 0.5, and the residual-scale prior has mean 0.2 MPa.

Table S7: Priors chosen for the hierarchical Bayesian model.

| Quantity | Prior | Interpretation |
|---|---|---|
| $\log m_{\mu_1}$ | $\mathcal{N}(\log 2, 1.25^2)$ | Population log-median of $\mu_1$ |
| $\log m_{\alpha_1}$ | $\mathcal{N}(\log 4, 0.80^2)$ | Population log-median of $\alpha_1$ |
| $z_{\mu_1,V}, z_{\alpha_1,V}$ | $\mathcal{N}(0, 1)$ | Standardized non-centered valve offsets |
| $z_{\mu_1,i}, z_{\alpha_1,i}$ | $\mathcal{N}(0, 1)$ | Standardized non-centered specimen offsets |
| $\sigma_{\mu_1,V}, \sigma_{\alpha_1,V}$ | Exponential($rate$ = 2) | Between-valve log-parameter standard deviations (dimensionless) |
| $\sigma_{\mu_1,S}, \sigma_{\alpha_1,S}$ | Exponential($rate$ = 2) | Between-specimen log-parameter standard deviations (dimensionless) |
| $\beta_{\mu_1,\log A}, \beta_{\alpha_1,\log A}$ | $\mathcal{N}(0, 0.25^2)$ | Regularized slopes, the change in log material parameter per 1 standard deviation of (log(Area)) (dimensionless) |
| $\sigma_{m,V}$ | Exponential($rate$ = 5MPa$^{-1}$) | Valve-specific residual stress standard deviation (MPa) |

The posterior combines the observed stress data $\mathbf{y}$ with the prior distribution,

$$p(\boldsymbol{\Theta} \mid \mathbf{y}) = \frac{p(\mathbf{y} \mid \boldsymbol{\Theta})p(\boldsymbol{\Theta})}{p(\mathbf{y})} \propto \left[\prod_i \prod_j p\{y_{ij} \mid f(\lambda_{ij}; \boldsymbol{\theta}_i), \sigma_{m,V[i]}\}\right] p(\boldsymbol{\Theta}), \tag{S34}$$

where $\boldsymbol{\Theta}$ contains the population coefficients, hierarchy scales, valve and specimen offsets, and residual stress scales.

#### *S4.1. Sampling and Diagnostics*

Posterior sampling was performed in Python 3.12.13 using PyMC 5.26.1 with the Nutpie 0.16.11 implementation of the No-U-Turn Sampler (NUTS). Model expressions and automatic differentiation were handled by PyTensor 2.35.1, and posterior summaries and convergence diagnostics were calculated using ArviZ 0.23.4. Four independent chains were sampled, each with 8,000 tuning iterations followed by 8,000 retained draws, yielding 32,000 retained posterior draws. The target acceptance probability was 0.97.

We assessed sampling using rank-normalized $\hat{R}$, effective sample size (ESS), divergences, and trace plots. Values of $\hat{R}$ close to one indicate agreement between within- and between-chain variation, with $\hat{R} < 1.01$ used as a convergence target. ESS estimates the equivalent number of independent draws for a posterior summary. Larger ESS reduces Monte Carlo uncertainty in that summary. The final run had zero divergences, a maximum $\hat{R}$ of 1.00, and a minimum bulk ESS of 1,603. The hierarchy-scale traces and marginal densities showed agreement among chains (Figure S3).

Figure S4 shows that the bulk and tail ESS increased with the number of retained draws. Bulk ESS describes sampling efficiency for central posterior summaries, while tail ESS is relevant to tail probabilities and interval endpoints. The approximately linear growth indicates that additional draws continued to increase the effective sample size. Table S8 summarizes the population medians and area slopes. A 95% highest-density interval (HDI) contains 95% of posterior probability in the region of highest density. Sampling diagnostics assess numerical exploration of the posterior and should be considered alongside checks of the fitted stress-stretch responses.

##### *S4.1.1. Area-Matched Specimen Posteriors and Fits*

To illustrate specimen-specific parameter inference while controlling for the model's only fixed covariate, we selected one specimen from each leaflet whose measured cross-sectional areas were as similar as possible. Among all mechanically tested specimens, we identified the posterior, anterior, and septal specimens that minimized the range in cross-sectional area, without

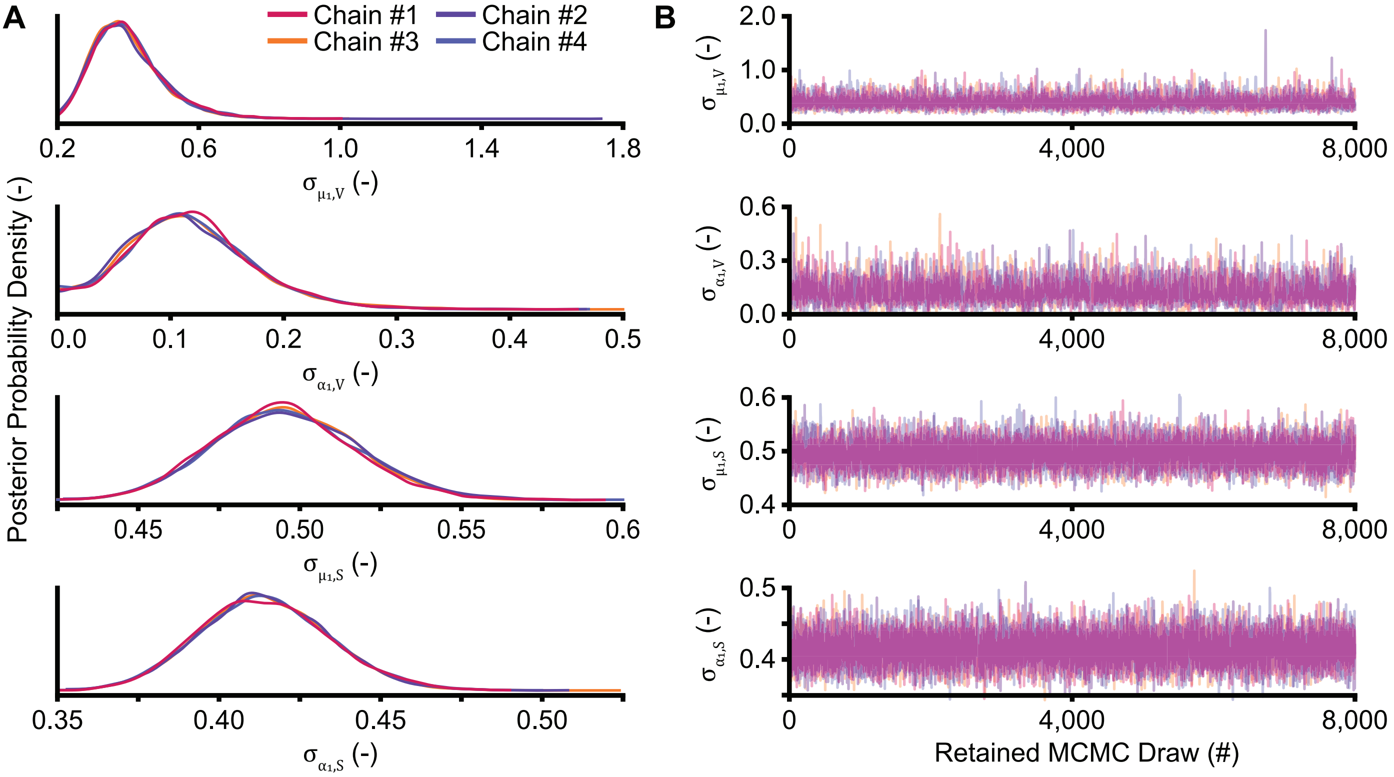


Figure S3: Hierarchy-scale diagnostics for the final model, with 8,000 retained draws per chain after 8,000 tuning iterations in each of four chains. **(A)** Each chain's posterior probability density closely agrees with the others for each hierarchy-scale parameter, suggesting the same posterior distribution is sampled. **(B)** Across the 8,000 retained posterior draws per chain, each chain's sampling band appears stationary with no sustained trends or shifts for each hierarchy-scale parameter. One hierarchy-scale parameter is shown per row, where $\sigma_{k,V}$ describes between-valve variation and $\sigma_{k,S}$ describes remaining between-specimen variation for each $k = [\mu_1, \alpha_1]$ parameter of the final Ogden-1 model. The hierarchy scales describe standard deviations of log material parameters and are dimensionless.

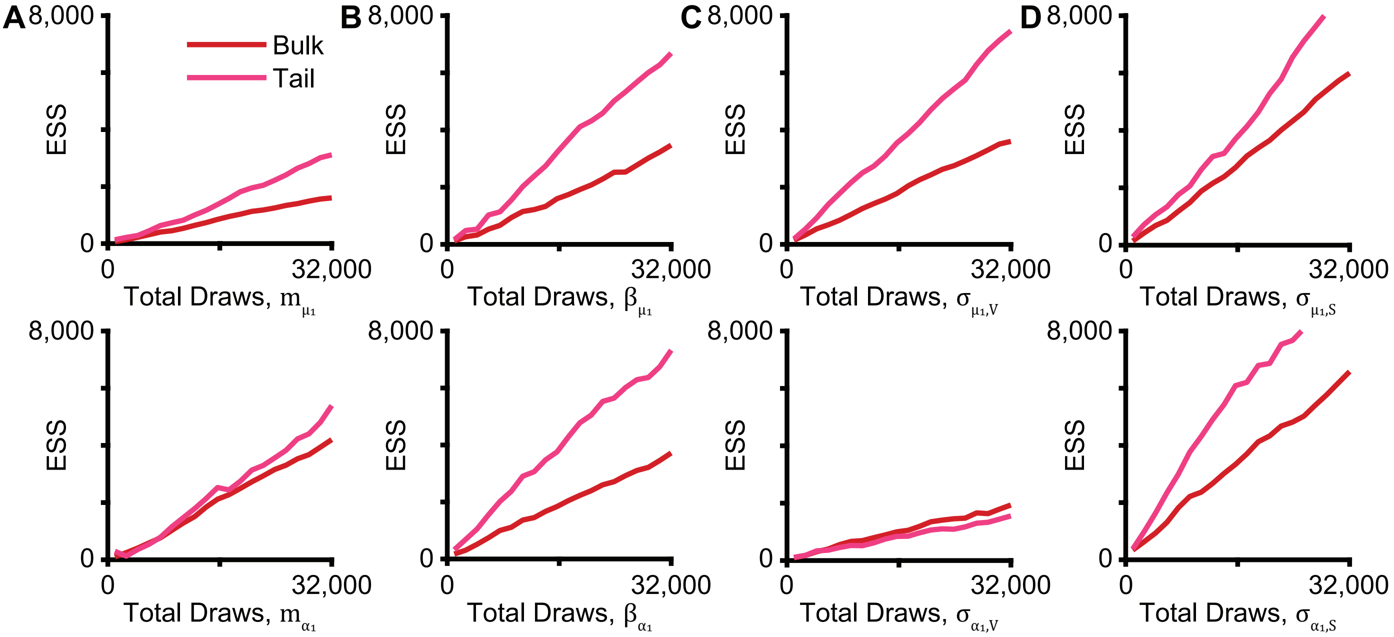


Figure S4: The effective sample size (ESS) sustains approximately linear growth for bulk and tail posterior summaries across 32,000 total retained draws in the final model. The top row shows the ESS progression for $\mu_1$ and bottom row for $\alpha_1$, the two parameters of the Ogden-1 model, for the posterior quantity of the **(A)** population medians, **(B)** log(Area) coefficients, **(C)** valve-level hierarchy scale, and the **(D)** specimen-level hierarchy scale.

Table S8: Summaries of the principal posteriors used in the final model. HDI = highest-density interval.

| Quantity | Mean | 95% HDI Lower | 95% HDI Upper |
|---|---|---|---|
| $m_{\mu_1}$ | 10.310 | 7.864 | 12.792 |
| $m_{\alpha_1}$ | 50.623 | 45.394 | 55.095 |
| $\beta_{\mu_1,\log A}$ | −0.344 | −0.411 | −0.279 |
| $\beta_{\alpha_1,\log A}$ | −0.206 | −0.261 | −0.150 |

restricting insertion site or number of tests. The selected specimens had areas of 0.303695 (CTL1 Posterior Near-Annulus #3), 0.303614 (HF3 Anterior Free Edge #2), and 0.303532 mm$^2$ (HF2 Septal Belly #5), respectively. Because log-transformed Area was the only fixed predictor in the final model, this matching limits differences attributable to Area and highlights differences associated with the specimens' observed mechanical responses and inferred valve- and specimen-level effects.

Figure S5 compares the specimen-specific marginal and joint posteriors with the population-level posterior evaluated at the mean area of the three specimens, $A$ = 0.303614 mm$^2$. The specimen-specific posteriors condition on each specimen's observed stress-stretch data and its inferred valve and specimen effects. The population-level posterior was evaluated at the matched area with both random effects set to zero. Although the specimens had nearly identical areas, their inferred material parameters differed appreciably, illustrating the residual mechanical heterogeneity represented by the hierarchical model after accounting for Area.

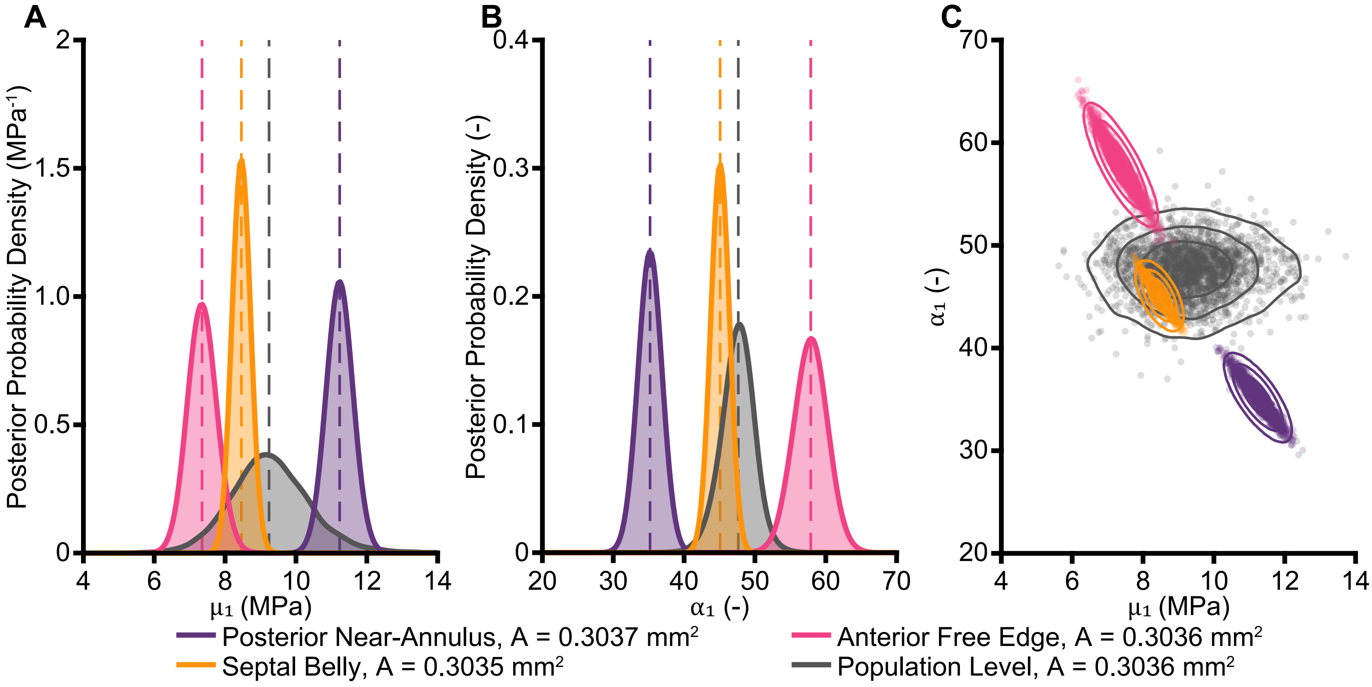


Figure S5: Area-matched specimen-specific and population-level Ogden-1 parameter posteriors. Panels **(A)** and **(B)** shows the marginal posterior probability densities of $\mu_1$ and $\alpha_1$, respectively. Colored distributions represent one posterior near-annulus specimen ($A$ = 0.303695 mm$^2$), one anterior free-edge specimen ($A$ = 0.303614 mm$^2$), and one septal belly specimen ($A$ = 0.303532 mm$^2$). The gray distribution is the population-level posterior evaluated at the mean area of the three specimens, $A$ = 0.303614 mm$^2$, with the valve- and specimen-level effects set to zero. Dashed vertical lines indicate posterior medians. Panel **(C)** shows the corresponding joint $(\mu_1, \alpha_1)$ posteriors. Translucent points show posterior draws and contours enclose approximately 50%, 80%, and 95% highest-density regions. Because Area is nearly identical among the three specimens, their posterior differences principally reflect their observed mechanical responses and inferred valve- and specimen-level effects rather than the fixed Area effect.

Figure S6 shows the corresponding stress-stretch fits predicted from the model. The pooled root-mean-square errors were 0.067 MPa for the posterior specimen, 0.096 MPa for the anterior specimen, and 0.098 MPa for the septal specimen. These errors were small relative to the approximately 3-MPa maximum measured stresses and indicate that the distinct specimen-specific posteriors were not caused by obvious model-fit failures. The specimens were selected for similarity in Area and representation of all three leaflets, rather than for having the lowest fitting errors.

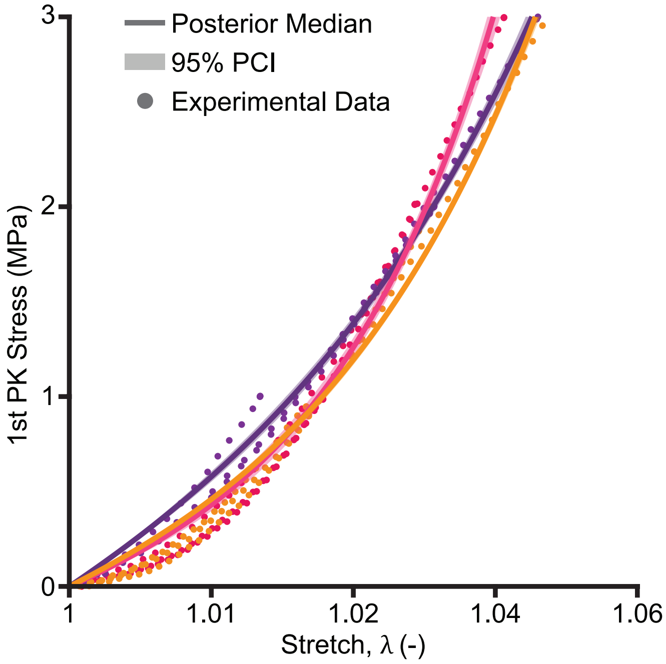


Figure S6: Experimental stress-stretch data and specimen-specific hierarchical Bayesian fits for the three area-matched chordae shown in Figure S5. Solid lines show the posterior-median latent Ogden-1 responses and shaded regions show pointwise 95% posterior intervals for the latent constitutive responses over the experimental observations (shown as circles). Colors identify the posterior near-annulus, anterior free-edge, and septal belly specimens. The posterior intervals propagate uncertainty in the specimen-specific material parameters but exclude Gaussian residual stress variation. All specimens had nearly identical cross-sectional areas of approximately 0.304 mm$^2$.

#### *S4.2. Population-Level Predictions*

To obtain a population-level response at a specified area $A$, we evaluated posterior draws with the valve and specimen random effects set to zero. For draw $s$, the material parameters are

$$\theta_k^{(s)}(A) = \exp\left[\log m_k^{(s)} + \beta_{k,\log A}^{(s)} \frac{\log A - \overline{\log A}}{s_{\log A}}\right]. \tag{S35}$$

These draws retain uncertainty in the population coefficients while excluding variation among individual valves and specimens. Area is held fixed. Evaluating Eq. S30 for each joint parameter draw gives a distribution of population-level constitutive responses. We retain $\mu_1$ and $\alpha_1$ from the same posterior draw to preserve their posterior dependence. Pointwise 95% posterior intervals for these curves exclude variation among new valves and specimens, uncertainty in area, and observation noise. Setting random effects to zero therefore does not remove uncertainty in the population-level response.

##### *S4.2.1. Prediction for a New Chorda from a New Valve*

For a new chorda $i^*$ from an unobserved valve $V^*$, predictions at fixed area $A^*$ additionally require fresh valve and specimen effects. For each posterior draw,

$$\theta_k^{*(s)}(A^*) = \exp\left[\log m_k^{(s)} + \beta_{k,\log A}^{(s)} \frac{\log A^* - \overline{\log A}}{s_{\log A}} + \sigma_{k,V}^{(s)} z_{k,V^*}^{(s)} + \sigma_{k,S}^{(s)} z_{k,i^*}^{(s)}\right], \qquad z_{k,V^*}^{(s)}, z_{k,i^*}^{(s)} \sim \mathcal{N}(0,1). \tag{S36}$$

This prediction includes posterior uncertainty in the population coefficients and hierarchy scales, together with variability among valves and specimens. It conditions on the specified area and excludes residual stress error. For multiple chordae belonging to the same simulated valve, the valve offset for each material parameter is shared, whereas each chorda receives its own specimen offset. The corresponding assigned area must also be used when converting nominal stress to axial force. These predictions therefore describe variability in the latent material parameters. Observation-level stress predictions would additionally require a specified residual stress scale and sampling of residual stress error.

Figure S7 compares the population-level and new-chorda parameter distributions at three fixed areas. The comparison at the pooled median area isolates the additional dispersion associated with valve and specimen variability, while comparisons across areas

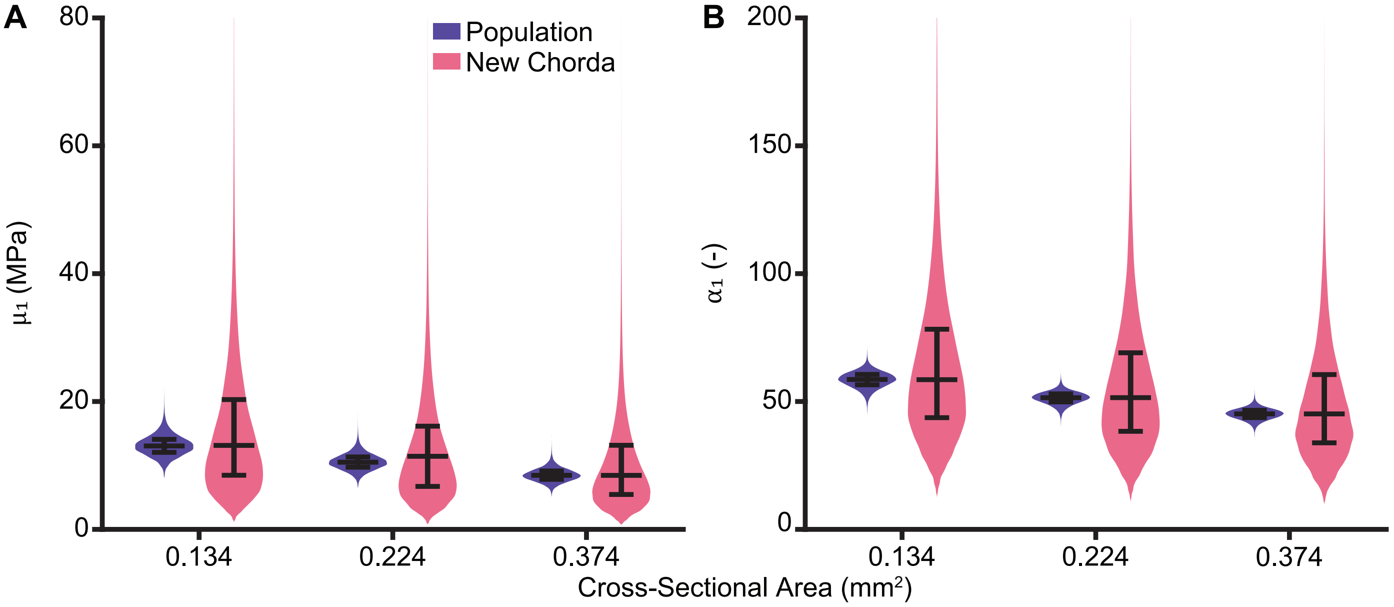


Figure S7: Population-level and new-chorda conditional distributions of the Ogden-1 parameters at fixed cross-sectional areas. Posterior distributions of **(A)** $\mu_1$ and **(B)** $\alpha_1$ were evaluated at the overall 25th-percentile $A_{25\%} = 0.134$ mm$^2$, median $A_{50\%} = 0.224$ mm$^2$, and 75th-percentile $A_{75\%} = 0.374$ mm$^2$ chordal cross-sectional areas. Population-level distributions were calculated with the valve- and specimen-level effects set to zero. New-chorda distributions additionally include newly sampled valve- and specimen-level effects at each joint posterior draw, representing a new chorda from a previously unseen valve. Cross-sectional area was held fixed within each comparison, so the additional dispersion reflects estimated between-valve and between-specimen heterogeneity rather than variation in area. Bars indicate the median and span the interquartile range.

illustrate the area dependence of the material parameters. Thus, at each fixed Area, the population-level distributions represent the conditional marginal posteriors $p(\mu_1 \mid A, \text{data})$ and $p(\alpha_1 \mid A, \text{data})$, whereas the new-chorda distributions represent $p(\mu_1^* \mid A^*, \text{data})$ and $p(\alpha_1^* \mid A^*, \text{data})$ after sampling fresh valve- and specimen-level effects.

#### *S4.2.2. Representative Anatomical Assignments*

When individual chordal areas are unavailable, the measured anatomical distributions can supply representative areas for model assignment. Table S9 gives median and interquartile areas for the 254 mechanically tested specimens with recorded insertion sites, combining control and heart failure groups. Of the 281 excised specimens, 16 lacked usable mechanical data and 11 lacked insertion-site information. The latter 11 specimens were included in the Bayesian fit because area was its only covariate, but were excluded from leaflet-by-insertion assignments.

Table S9: Chordae Tendineae median cross-sectional area (mm$^2$) and interquartile range (IQR) by leaflet and insertion site. 254 specimens are included, where **n** is the number of specimens per group, and both control and heart failure groups are combined.

| Leaflet | Insertion Site | n = | Median Cross-Sectional Area (mm$^2$) | IQR [25%–75%] |
|---|---|---|---|---|
| Posterior | Near-Annulus | 18 | 0.176 | 0.120–0.300 |
| | Belly | 28 | 0.234 | 0.119–0.365 |
| | Free Edge | 32 | 0.255 | 0.153–0.353 |
| Anterior | Near-Annulus | 21 | 0.372 | 0.276–0.726 |
| | Belly | 33 | 0.454 | 0.249–0.644 |
| | Free Edge | 33 | 0.213 | 0.134–0.340 |
| Septal | Near-Annulus | 20 | 0.135 | 0.087–0.201 |
| | Belly | 37 | 0.198 | 0.155–0.304 |
| | Free Edge | 32 | 0.132 | 0.101–0.222 |

We evaluated Eq. S35 at each group's median area. Leaflet and insertion site influence these predictions through the assigned areas and are not coefficients in the final regression. For deterministic simulations, we selected an actual joint posterior draw whose predicted curve approximated the pointwise posterior-median stress-stretch curve at the specified area. Thus, selecting a representative parameter pair is a separate step from generating the posterior distribution. The representative assignments are reported in Table 2, and their curves are compared with the experimental data in Figure 7. All representative parameter assignments, posterior summaries, and uncertainty intervals were obtained from the final fit with 8,000 tuning iterations and 8,000 retained draws per chain across four chains. Results from the initial fit are reported only for the sampling-sensitivity comparison in Section S4.3.

We repeated this procedure at each group's 25th- and 75th-percentile areas. Table S10 reports the resulting representative joint parameter pairs. These assignments describe changes with area and should not be interpreted as credible-interval endpoints for the parameters.

Table S10: Population-level Ogden-1 parameter predictions at the 25% and 75% quartiles of cross-sectional area within each leaflet-by-insertion cell.

| **Leaflet** | **Insertion** | $A_{25\%}$ (mm$^2$) | $\mu_{1,25\%}$ (MPa) | $\alpha_{1,25\%}$ (-) | $A_{75\%}$ (mm$^2$) | $\mu_{1,75\%}$ (MPa) | $\alpha_{1,75\%}$ (-) |
|---|---|---|---|---|---|---|---|
| | Near-Annulus | 0.120 | 13.473 | 60.348 | 0.300 | 9.261 | 47.708 |
| Posterior | Belly | 0.119 | 13.556 | 60.510 | 0.365 | 8.552 | 45.284 |
| | Free Edge | 0.153 | 12.210 | 56.742 | 0.353 | 8.549 | 45.964 |
| | Near-Annulus | 0.276 | 9.561 | 48.652 | 0.726 | 6.371 | 38.023 |
| Anterior | Belly | 0.249 | 9.971 | 49.973 | 0.644 | 6.728 | 39.109 |
| | Free Edge | 0.134 | 13.068 | 58.382 | 0.340 | 8.785 | 46.103 |
| | Near-Annulus | 0.087 | 15.448 | 65.719 | 0.201 | 10.787 | 53.052 |
| Septal | Belly | 0.155 | 12.143 | 56.479 | 0.304 | 9.212 | 47.543 |
| | Free Edge | 0.101 | 14.573 | 63.136 | 0.222 | 10.407 | 51.688 |

Table S11 reports representative assignments at median areas for all mechanically tested specimens and for groups defined by heart failure status, leaflet, or insertion site. These group labels are not predictors in the Bayesian model. Differences among the assignments arise from the selected group median areas.

Table S11: Population-level Ogden-1 parameters predicted using cross-sectional area pooled over all data or within one grouping factor, where **n** is the number of specimens included in the group.

| **Group** | **n =** | **Median Area (mm$^2$)** | $\mu_1$ **(MPa)** | $\alpha_1$ **(-)** |
|---|---|---|---|---|
| All | 265 | 0.224 | 10.367 | 51.569 |
| Control | 148 | 0.220 | 10.464 | 51.859 |
| Heart Failure | 117 | 0.236 | 10.222 | 50.733 |
| Posterior | 81 | 0.247 | 9.991 | 50.033 |
| Anterior | 92 | 0.343 | 8.754 | 46.000 |
| Septal | 92 | 0.165 | 11.905 | 55.605 |
| Near-Annulus | 59 | 0.207 | 10.657 | 52.743 |
| Belly | 98 | 0.248 | 9.989 | 50.027 |
| Free Edge | 97 | 0.195 | 11.128 | 53.117 |

Figure S8 compares the representative Bayesian responses with the least-squares fits to group median curves. The Bayesian responses were more compliant over the plotted range. The approaches differed in calibration range and statistical target. Grouped least-squares fits used 0–1 MPa and fitted a curve constructed by first taking the median across repeated tests within specimens and then across specimens. Bayesian fitting used all available loading data and represented variation among specimens and valves through the hierarchy. These differences may help explain the differing parameter estimates, but their individual contributions were not isolated.

#### *S4.3. Sensitivity of Representative Ogden Parameters to MCMC Sampling*

We first used an initial fit with four chains, 2,000 tuning iterations and 2,000 retained draws per chain (8,000 total retained posterior draws), and a target acceptance probability of 0.92. To evaluate the sensitivity of the predicted parameters to the amount of posterior sampling and sampler control, we repeated the otherwise identical model using four chains, 8,000 tuning iterations and 8,000 retained draws per chain (32,000 total retained posterior draws), and a target acceptance probability of 0.97. The measured data, model structure, priors, log-area standardization, cell-specific median and quartile areas, and representative joint-draw selection procedure were unchanged between fits.

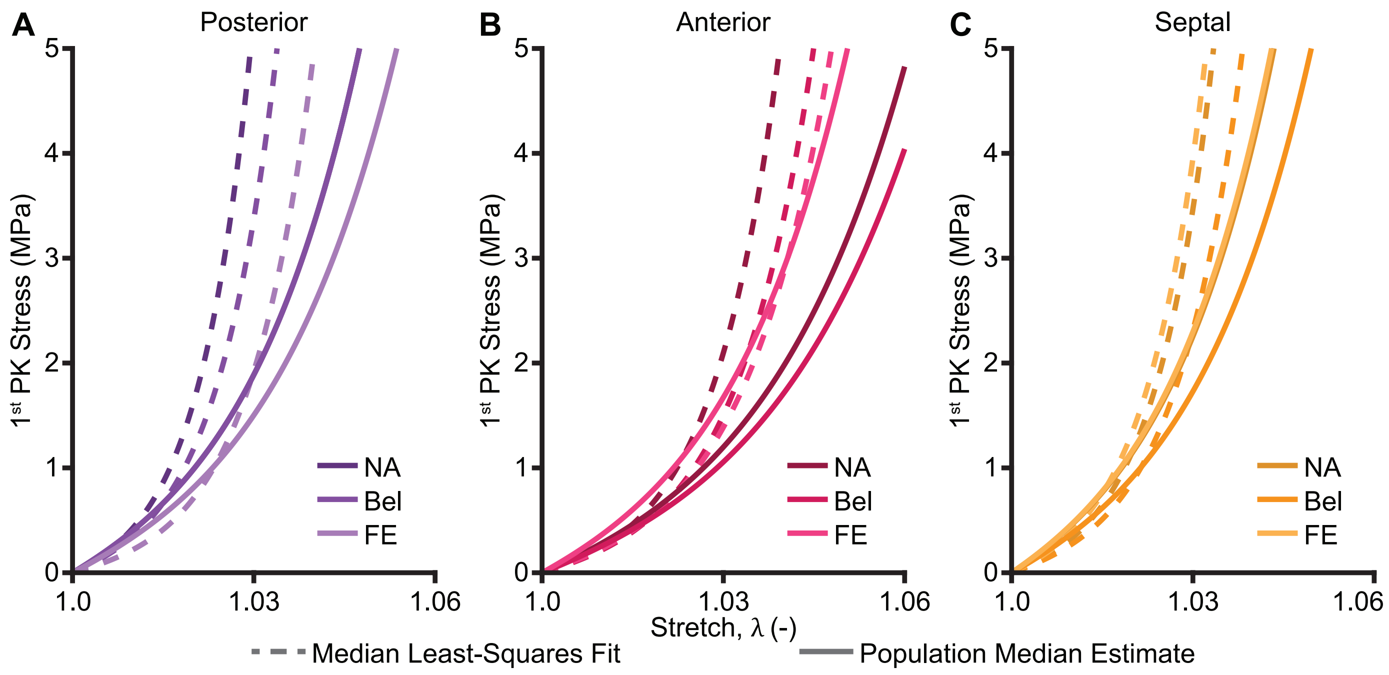


Figure S8: Comparison of Ogden-1 stress-stretch responses by leaflet and insertion site. Dashed lines show separate least-squares fits to each group's median stress-stretch curve. Solid lines show representative predictions from the hierarchical Bayesian model evaluated at each group's median cross-sectional area. Panels show the **(A)** posterior, **(B)** anterior, and **(C)** septal leaflets. Least-squares fits used data from 0–1 MPa and were extrapolated beyond 1 MPa, whereas the Bayesian model used all available loading data. The Bayesian responses were more compliant over the plotted range.

For each leaflet-by-insertion cell, we evaluated the low-sampling (LS) and high-sampling (HS) representative Ogden-1 curves over $1 < \lambda \leq 1.06$. We defined their pointwise relative difference and parameter difference as,

$$e(\lambda) = 100\,\frac{P_{\mathrm{LS}}(\lambda) - P_{\mathrm{HS}}(\lambda)}{P_{\mathrm{HS}}(\lambda)}, \qquad \Delta\mu_1 = 100\frac{\mu_{1,\mathrm{LS}} - \mu_{1,\mathrm{HS}}}{\mu_{1,\mathrm{HS}}}, \qquad \Delta\alpha_1 = 100\frac{\alpha_{1,\mathrm{LS}} - \alpha_{1,\mathrm{HS}}}{\alpha_{1,\mathrm{HS}}}. \tag{S37}$$

where $P$ is the predicted first Piola-Kirchhoff stress. The undeformed point $\lambda = 1$ was excluded from the relative calculation because both models predict zero stress there.

Increasing the number of retained posterior draws and the target acceptance probability produced only minor changes in the representative properties. Across all median and quartile assignments, the absolute change in $\mu_1$ was no greater than 2.79%, while the absolute change in $\alpha_1$ was no greater than 0.64%. Every low-sampling parameter value remained within the corresponding 95% posterior interval from the high-sampling fit. The maximum relative difference between corresponding stress-stretch curves was 2.79%. Curve-wise RMSE was at most 0.113 MPa, and the largest absolute pointwise stress difference was 0.263 MPa. At $\lambda = 1.06$, the relative differences ranged from 0.34% to 2.11%. Thus, the representative median- and quartile-area constitutive responses were insensitive to the increased posterior sampling and target acceptance probability. The high-sampling fit was used for the final posterior summaries and uncertainty intervals reported in this study.

Table S12: Sensitivity of representative population-level Ogden-1 properties to MCMC sampling comparing the initial and final hierarchical Bayesian fits at each cell's 25th-percentile, median, and 75th-percentile cross-sectional area. The low-sampling (LS) fit used four chains with 2,000 tuning and 2,000 retained iterations per chain and a target acceptance probability of 0.92. Parameter differences and constitutive-curve errors are calculated relative to the high-sampling fit, which used 8,000 tuning and 8,000 retained iterations per chain and a target acceptance probability of 0.97. Stress errors were evaluated over ($1 < \lambda \leq 1.06$) and metrics were calculated using the full-precision parameter exports.

| **Leaflet** | **Insertion** | **Quartile** | $A$ **(mm$^2$)** | $\mu_{1,\text{LS}}$ **(MPa)** | $\alpha_{1,\text{LS}}$ **(-)** | $\Delta\mu_1$ **(%)** | $\Delta\alpha_1$ **(%)** | **Stress RMSE (MPa)** | **Max.** $\lvert e(\lambda)\rvert$ **(%)** |
|---|---|---|---|---|---|---|---|---|---|
| | | 25% | 0.120 | 13.709 | 60.242 | 1.75 | -0.18 | 0.074 | 1.75 |
| Posterior | Near-Annulus | Median | 0.176 | 11.623 | 54.868 | 0.22 | 0.44 | 0.039 | 1.21 |
| | | 75% | 0.300 | 9.355 | 47.795 | 1.02 | 0.18 | 0.032 | 1.37 |
| | | 25% | 0.119 | 13.792 | 60.447 | 1.74 | -0.10 | 0.083 | 1.74 |
| Posterior | Belly | Median | 0.234 | 10.387 | 50.774 | 1.32 | -0.10 | 0.035 | 1.32 |
| | | 75% | 0.365 | 8.683 | 45.190 | 1.53 | -0.21 | 0.026 | 1.53 |
| | | 25% | 0.153 | 12.344 | 56.769 | 1.09 | 0.05 | 0.051 | 1.21 |
| Posterior | Free edge | Median | 0.255 | 10.022 | 49.734 | 0.72 | 0.06 | 0.023 | 0.84 |
| | | 75% | 0.353 | 8.647 | 45.938 | 1.15 | -0.06 | 0.023 | 1.15 |
| | | 25% | 0.276 | 9.685 | 48.760 | 1.30 | 0.22 | 0.042 | 1.73 |
| Anterior | Near-Annulus | Median | 0.372 | 8.616 | 45.026 | 1.55 | -0.27 | 0.025 | 1.55 |
| | | 75% | 0.726 | 6.495 | 38.036 | 1.95 | 0.03 | 0.025 | 1.99 |
| | | 25% | 0.249 | 10.133 | 50.050 | 1.62 | 0.15 | 0.052 | 1.93 |
| Anterior | Belly | Median | 0.454 | 7.927 | 42.776 | 2.74 | -0.38 | 0.040 | 2.73 |
| | | 75% | 0.644 | 6.782 | 39.340 | 0.79 | 0.59 | 0.020 | 1.63 |
| | | 25% | 0.134 | 13.091 | 58.615 | 0.18 | 0.40 | 0.048 | 1.16 |
| Anterior | Free edge | Median | 0.213 | 10.815 | 51.970 | 1.52 | -0.56 | 0.020 | 1.52 |
| | | 75% | 0.340 | 8.817 | 46.321 | 0.37 | 0.47 | 0.023 | 1.20 |
| | | 25% | 0.087 | 15.847 | 65.425 | 2.59 | -0.45 | 0.113 | 2.59 |
| Septal | Near-Annulus | Median | 0.135 | 13.079 | 58.584 | 0.17 | 0.39 | 0.047 | 1.13 |
| | | 75% | 0.201 | 11.084 | 52.713 | 2.75 | -0.64 | 0.055 | 2.75 |
| | | 25% | 0.155 | 12.264 | 56.560 | 0.99 | 0.14 | 0.054 | 1.33 |
| Septal | Belly | Median | 0.198 | 11.159 | 52.920 | 2.79 | -0.55 | 0.062 | 2.79 |
| | | 75% | 0.304 | 9.305 | 47.647 | 1.02 | 0.22 | 0.032 | 1.42 |
| | | 25% | 0.101 | 14.795 | 62.897 | 1.52 | -0.38 | 0.044 | 1.52 |
| Septal | Free edge | Median | 0.132 | 13.202 | 58.893 | -0.18 | 0.32 | 0.024 | 0.61 |
| | | 75% | 0.222 | 10.624 | 51.437 | 2.08 | -0.48 | 0.040 | 2.08 |

### S5. Additional Results

#### *S5.1. Chordal Insertion Density*

Raw chordal counts for each mesh cell are given in Supplement Table S13. We divide the raw count by the number of cells to compute an average chordal number per cell in Supplement Table S14. Note that this is not a density, as cell area is not taken into account. Cell areas and densities are available with the raw data.

Table S13: Total number of chordal insertions identified in each mesh cell, grouped by heart failure (CTL = control, HF = heart failure), leaflet (Post = posterior, Ant = anterior, Sept = septal), radial position (NA = near-annulus, Bel = belly, FE = free edge), and circumferential position (L = left, C = central, R = right).

| | **Total** | **Heart Failure** | | **Leaflet** | | | **Radial Position** | | | **Circumferential Position** | | |
|---|---|---|---|---|---|---|---|---|---|---|---|---|
| | | CTL | HF | Post | Ant | Sept | NA | Bel | FE | L | C | R |
| **Total** | 5,710 | 3,255 | 2,455 | 1,649 | 2,272 | 1,789 | 627 | 2,025 | 3,058 | 1,702 | 2,534 | 1,474 |
| CTL | | - | - | 959 | 1,325 | 971 | 413 | 1,283 | 1,559 | 986 | 1,437 | 832 |
| HF | | | - | 690 | 947 | 818 | 214 | 742 | 1,499 | 716 | 1,097 | 642 |
| Post | | | | - | - | - | 191 | 570 | 888 | 473 | 713 | 463 |
| Ant | | | | | - | - | 241 | 827 | 1,204 | 739 | 1,000 | 533 |
| Sept | | | | | | - | 195 | 628 | 966 | 490 | 821 | 478 |
| NA | | | | | | | - | - | - | 179 | 286 | 162 |
| Bel | | | | | | | | - | - | 636 | 818 | 571 |
| FE | | | | | | | | | - | 887 | 1,430 | 741 |

Table S14: Average number of chordal insertions per mesh cell, calculated as the total number of insertions divided by the corresponding number of cells, grouped by heart failure (CTL = control, HF = heart failure), leaflet (Post = posterior, Ant = anterior, Sept = septal), radial position (NA = near-annulus, Bel = belly, FE = free edge), and circumferential position (L = left, C = central, R = right). These values represent average insertion counts per cell rather than insertion densities normalized by area.

| | **Total** | **Heart Failure** | | **Leaflet** | | | **Radial Position** | | | **Circumferential Position** | | |
|---|---|---|---|---|---|---|---|---|---|---|---|---|
| | | CTL | HF | Post | Ant | Sept | NA | Bel | FE | L | C | R |
| **Total** | 6.41 | 6.35 | 6.49 | 5.23 | 6.64 | 7.65 | 2.11 | 6.82 | 10.30 | 5.73 | 8.53 | 4.96 |
| CTL | | - | - | 5.33 | 6.69 | 7.19 | 2.42 | 7.50 | 9.12 | 5.77 | 8.40 | 4.87 |
| HF | | | - | 5.11 | 6.58 | 8.26 | 1.70 | 5.89 | 11.90 | 5.68 | 8.71 | 5.10 |
| Post | | | | - | - | - | 1.82 | 5.43 | 8.46 | 4.50 | 6.79 | 4.41 |
| Ant | | | | | - | - | 2.11 | 7.25 | 10.56 | 6.48 | 8.77 | 4.68 |
| Sept | | | | | | - | 2.50 | 8.05 | 12.38 | 6.28 | 10.53 | 6.13 |
| NA | | | | | | | - | - | - | 1.81 | 2.89 | 1.64 |
| Bel | | | | | | | | - | - | 6.42 | 8.26 | 5.77 |
| FE | | | | | | | | | - | 8.96 | 14.44 | 7.48 |

Figure S9 shows observed chordal insertion densities grouped by heart failure, leaflet, radial, and circumferential position. Conditional chordal insertion rates were lower in the anterior leaflet than the septal or posterior leaflets ($p < 0.001$), but not significantly different between the septal and posterior leaflets ($p = 0.576$). Insertion rates increased from the annulus to the free-edge (all $p < 0.001$). Interestingly, cells labeled right had lower insertion rates than those labeled central ($p = 0.004$) or left ($p = 0.004$), but the left and central cells had similar insertion rates ($p = 0.818$). Because circumferential labels were assigned locally within each scallop, and adjacent scallops share internal boundaries, this finding should be interpreted as an asymmetry in the parameterized scallop coordinates rather than evidence of a global anatomical left-right asymmetry. The main effect of heart failure was not significant ($p = 0.057$), but its interaction with radial position was significant ($p < 0.001$). Conditional rates were lower in heart failure than in controls at the near-annulus ($p = 0.025$) and belly ($p = 0.004$), but not at the free edge ($p = 0.829$).

To view the interaction of the radial and circumferential position on chordal insertion density, we show the descriptive insertion density grouped spatially between radial and circumferential position in Figure S10. Since the interaction between radial and circumferential position were excluded from the final model, we do not make statistical inferences for these data.

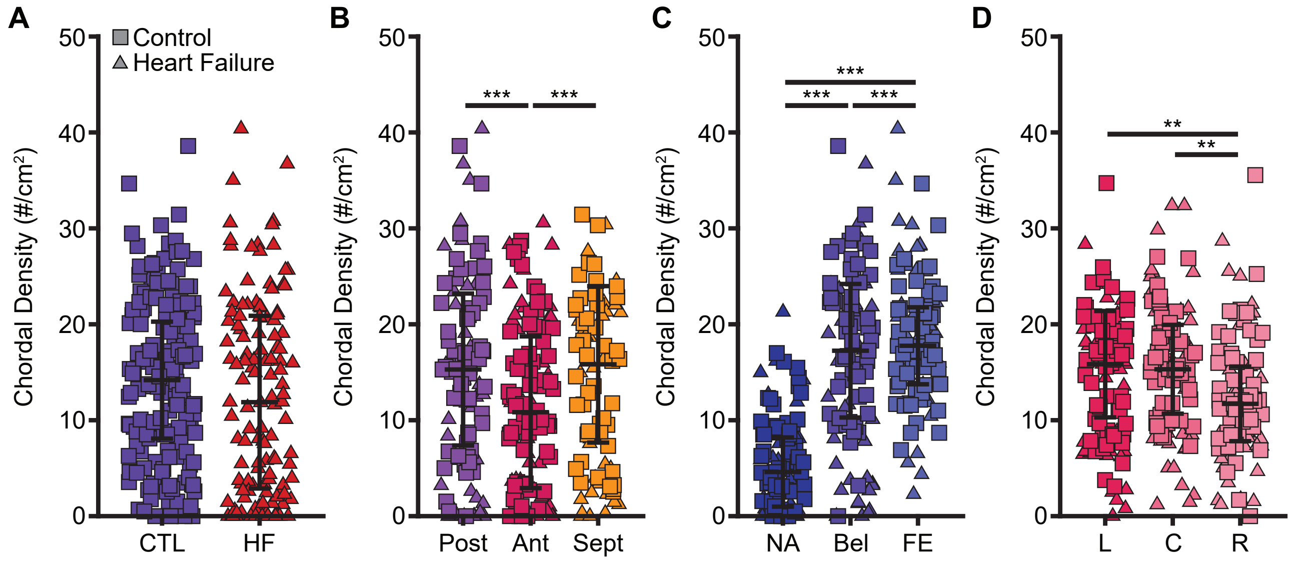


Figure S9: Chordal insertion densities and conditional insertion-rate comparisons by **(A)** heart failure, **(B)** leaflet, **(C)** insertion site (radial position), and **(D)** circumferential position. Each observation represents insertion count divided by cell area for one of 891 cells across 99 scallops. Summary bars indicate medians and interquartile range. Statistical comparisons refer to the conditional negative-binomial component, rather than overall expected density incorporating zero inflation. Asterisks indicate statistical significance (* $p<0.05$, ** $p<0.01$, *** $p<0.001$).

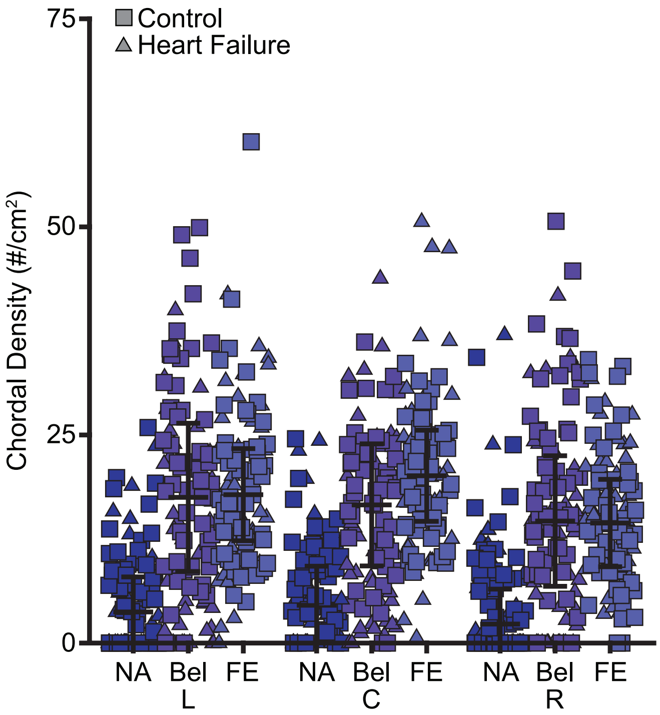


Figure S10: Chordae tendineae insertion density for all cells of the 3 × 3 grid for all leaflets and grouped across control and heart failure hearts. Across 891 cells, we group by radial (NA/Bel/FE) and then by circumferential position (L/C/R), where error bars indicate the median and interquartile range. No statistical comparisons between groups were performed.

*S5.2. Chordae Tendineae Cross-Sectional Area*

To investigate the spatial interaction on chordal cross-sectional area, we show the cross-sectional area grouped by insertion site and then by leaflet in Figure S11. As clearly shown, there are many significant differences in the median cross-sectional area for each leaflet-by-insertion position. We point the interested reader to the full data and R codes as described in the Data Availability Statement 8 to generate a full list of these results. In short, the anterior leaflet has larger chordae than the septal or posterior, while the septal free edge has the smallest chordae.

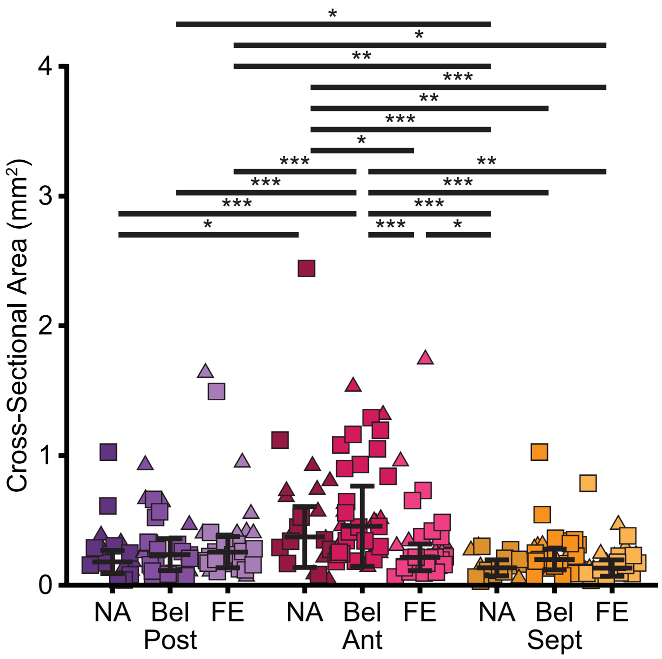


Figure S11: Chordae tendineae cross-sectional area is different for each leaflet-by-insertion site. Cross-sectional area is grouped by insertion site and then by leaflet for 270 specimens with valid insertion data. We show cross-sectional area for control and heart failure, where error bars indicate the median and interquartile range. Asterisks indicate statistical significance (* $p<0.05$, ** $p<0.01$, *** $p<0.001$).

*S5.3. Mechanical Results*

We computed four metrics in order to quantify the mechanical behavior of the tested specimens: toe stiffness, calf stiffness, transition stretch, and normalized hysteresis. Toe and calf tangent stiffnesses were calculated as the slopes of linear fits to the three stress-stretch observations nearest 0.025 and 1 MPa, respectively. All 689 calf stiffness observations in the primary mixed-effects models (Figures 5, 6B, S13) were evaluated at 1 MPa. Additional tangent stiffnesses for Supplementary Figure S15 were similarly calculated near 2 and 3 MPa for tests reaching those loading levels. Transition stretch was defined as the intersection of the toe and calf fitted lines. When automated point selection produced a nonphysical transition stretch, the stiffness-fitting regions were expanded or manually adjusted for 31 specimens. The complete point-selection and fitting procedure, including adjustments, is provided in the accompanying analysis code. Calf stiffness is presented and discussed in the main manuscript, see Section 3.3, while the remaining three are presented here.

In Figure S12, we show the toe stiffness for each test when grouped by heart failure, leaflet, and insertion site. We found that specimen cross-sectional area ($p < 0.001$), peak stress ($p < 0.001$), and the heart failure-by-leaflet interaction ($p < 0.001$) were significant. As with calf stiffness, the main effects of heart failure ($p = 0.744$), leaflet ($p = 0.115$), and insertion site ($p = 0.170$) were not significant after adjustment for area and peak stress. This suggests that the mechanical behavior at small stresses is driven primarily by the size of the specimen.

Calf stiffness showed a significant interaction between area, leaflet, and insertion site ($p = 0.005$). In Figure S13, we plot the calf stiffness as a function of the cross-sectional area for each leaflet-by-insertion cell. While the differences between the calf stiffness of specimens inserting into the posterior and anterior leaflet are negligible, we see noticeable differences in the septal leaflet (Figure S13C) when comparing at a given area. Calf stiffness can serve as an approximation for the stiffness of the specimen at high stretches. Here, near-annulus septal chordae behave stiffer than belly chordae at the same size, while belly chordae become more compliant as they increase in size. We tested whether the slopes differed at the quartile values of cross-sectional area, and found

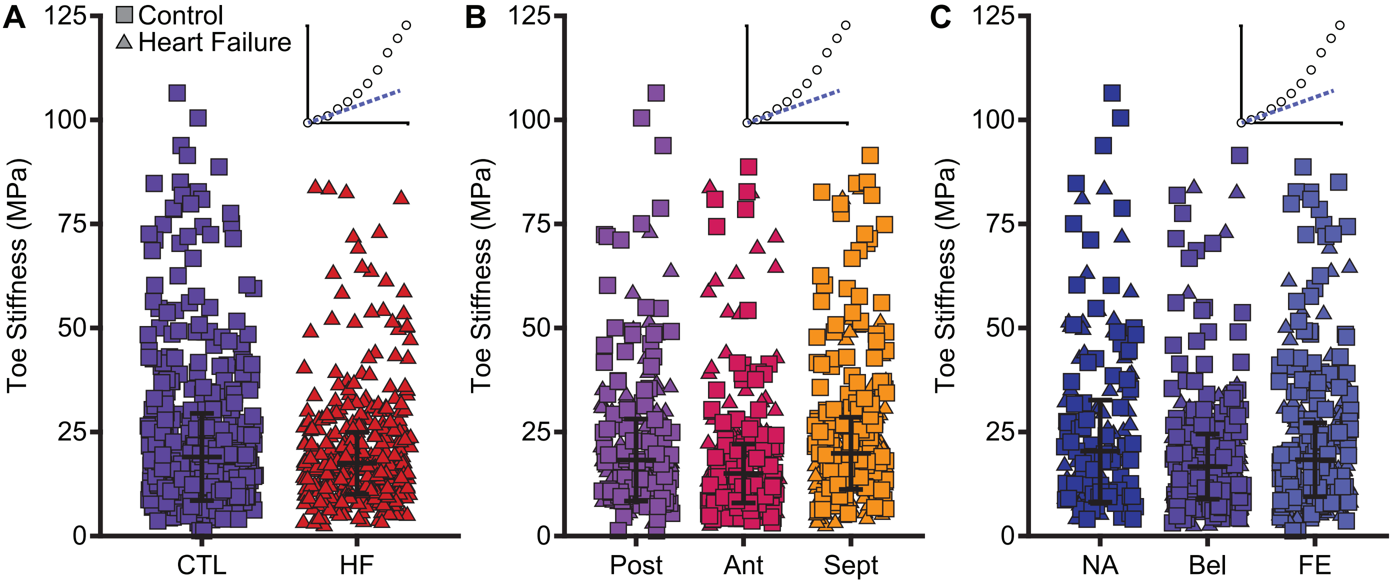


Figure S12: There is no statistical difference between chordal toe stiffness when specimens are grouped by **(A)** heart failure, **(B)** leaflet, or **(C)** insertion site after accounting for the cross-sectional area of each specimen. That is, pairwise comparisons between the toe stiffness for each group show no statistical differences when the cross-sectional area of the specimen is included as a covariate. Toe stiffness is computed as the tangent stiffness near 0 MPa (see insets), where each point is from one of 689 uniaxial tests, where error bars indicate the median and interquartile range. Only tests with a valid insertion site are shown.

that only the septal near-annulus chordae were stiffer than the septal belly chordae ($p = 0.025$) at a 75th percentile of area. Other comparisons at the evaluated area quartiles were not significant. Thus, while other chordae when grouped by leaflet-by-insertion become more compliant as their size increases, septal near-annulus chordae remain stiffer even at larger cross-sectional areas.

In addition to toe and calf stiffness, we also compute the transition stretch as the stretch at which the toe and calf tangent lines intersect. Toe stiffness can be interpreted as the stiffness at low stretch/stress during the soft response of the tissue before collagen fiber recruitment. Calf stiffness can be interpreted as the stiffness once the collagen fibers of the tissue engage and the mechanical response stiffens. Thus, the transition stretch can be interpreted as the point at which the material begins to stiffen and follow the classic non-linear J-shape. Figure S14 shows the transition stretch for each test when grouped by heart failure, leaflet, and insertion site. We found that specimen cross-sectional area ($p < 0.001$), peak stress ($p < 0.001$), and the heart failure-by-leaflet interaction ($p < 0.001$) were significant. Main effects of heart failure ($p = 0.843$), leaflet ($p = 0.554$), and insertion site ($p = 0.104$) were not significant. Within the septal leaflet, controls had lower transition stretch than heart-failure specimens ($p=0.043$). Within the anterior leaflet, near-annulus specimens had lower transition stretch than free-edge specimens ($p=0.014$).

We computed the calf stiffness at 1 MPa since all specimens tested reached at least 1 MPa. However, for those specimens tested to 2 or 3 MPa, we could have computed the calf stiffness at a larger stress. To understand how the tangent stiffness changes, we plot the value of the tangent stiffness as a function of where the tangent line is computed in Figure S15. Note that these fits are descriptive because they contain repeated tests. As expected for a nonlinear stress-stretch curve, we see that the tangent stiffness increases when moving along the loading curve. This increase is consistent across specimen groups and sizes. Thus, our choice to compute the calf stiffness at 1 MPa provided a simple way to take a snapshot of the mechanical stiffness at a specific stress between all tested specimens. We also fit linear models to all tangent stiffness points and for each grouping. While all models were statistically significant ($p < 0.001$), they only provided moderate ability for the stress at calculation to predict the tangent stiffness as given by the adjusted $R^2$. This is due to the fact that the stress-stretch curves are non-linear themselves, so their tangent stiffnesses also increase non-linearly. The linear fits for specimens grouped by chordal size provided the best predictability, which aligns with our prior findings on the importance of cross-sectional area in determining mechanical behavior. We conclude that the more complex hyperelastic models are needed to fully capture and compare the nonlinear mechanical behavior of the chordae tendineae. Lastly, we note that all specimens are tested from 0-1 MPa, but only some specimens reach 2 MPa, and even fewer reach 3 MPa. Therefore, there are more sample points at lower stresses than at higher stresses. Additionally, specimens are tested first to 1 MPa, then to 2 MPa, and then to 3 MPa, so order effects cannot be ruled out. To this end, we investigate order effects below.

We computed the hysteresis for each test as the area enclosed by the loading and unloading branches of the stress-stretch curve. The loading and unloading energy densities were calculated by trapezoidal integration of the $1^{st}$ Piola-Kirchhoff stress with respect to stretch, ($W_{\text{load}} = \int P_{\text{load}} d\lambda$) and ($W_{\text{unload}} = \int P_{\text{unload}} d\lambda$), respectively. The dissipated energy density was defined as the difference between the loading and returned energies ($W_{\text{diss}} = W_{\text{load}} - W_{\text{unload}}$). Since specimens are tested to different peak stresses and required larger loading energy, we normalized the hysteresis by the loading energy density. Thus, normalized hysteresis was defined as ($H_{\text{norm}} = W_{\text{diss}}/W_{\text{load}}$), representing the fraction of the mechanical energy applied during loading that was dissipated over

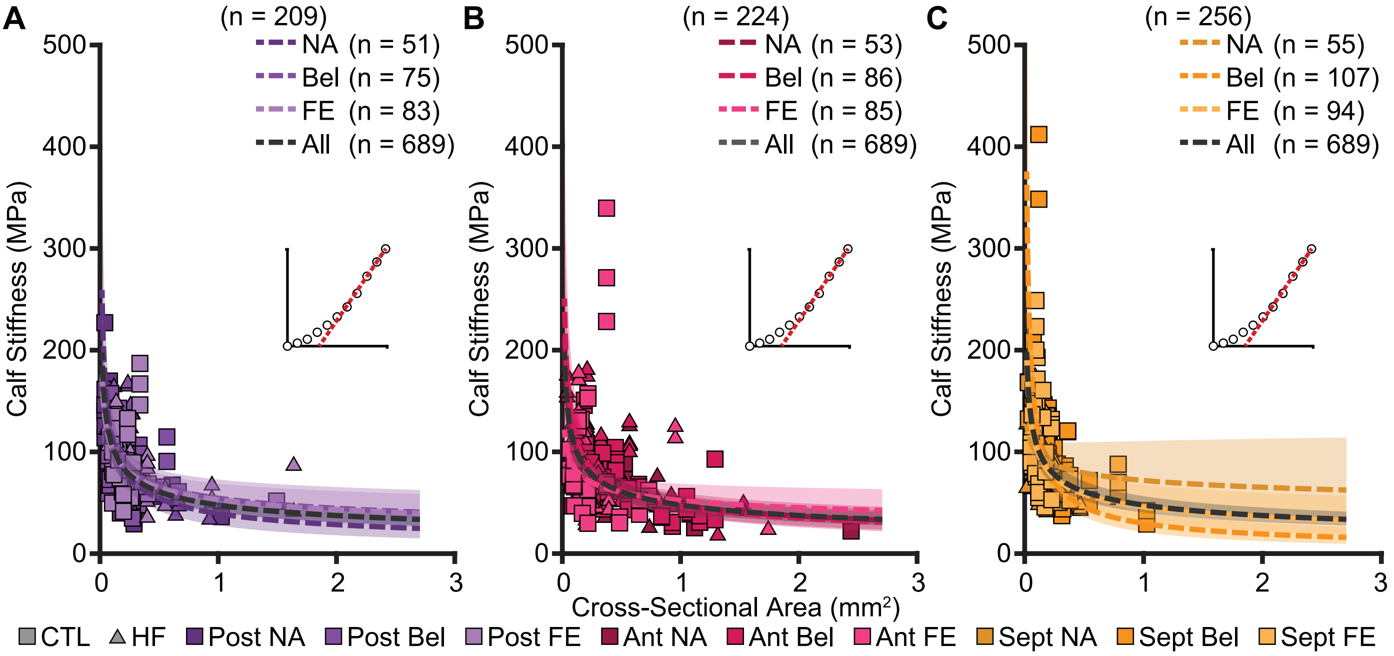


Figure S13: Calf stiffness of chordae inserting into the septal near-annulus remains larger at larger cross-sectional areas. In contrast, the calf stiffness of septal belly chordae becomes lower at larger areas. Here, calf stiffness is correlated to cross-sectional area for each insertion site in the **(A)** posterior, **(B)** anterior, and **(C)** septal leaflet. Calf stiffness is calculated as the tangent stiffness near 1 MPa for **n** of 689 total uniaxial tests, see insets. The dashed line depicts the correlation estimated by the linear mixed-effects model between the calf stiffness for each leaflet-by-insertion site group and cross-sectional area, with shading showing the 95% confidence interval.

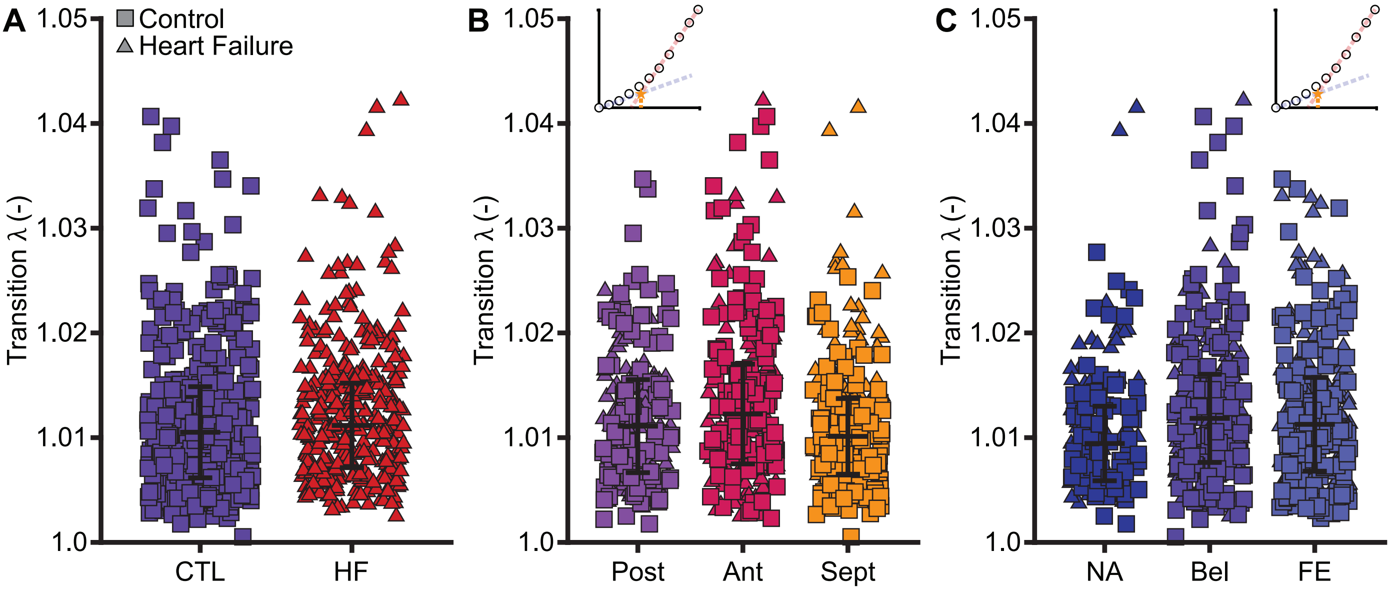


Figure S14: Overall pairwise comparisons showed no statistical difference between transition stretch when specimens are grouped by **(A)** heart failure, **(B)** leaflet, or **(C)** insertion site after accounting for the cross-sectional area of each specimen. Transition stretch is computed as the stretch at the intersection of the toe and calf stiffness (see insets), where each point is from one of 689 uniaxial tests, where error bars indicate the median and interquartile range. Only tests with a valid insertion site are shown.

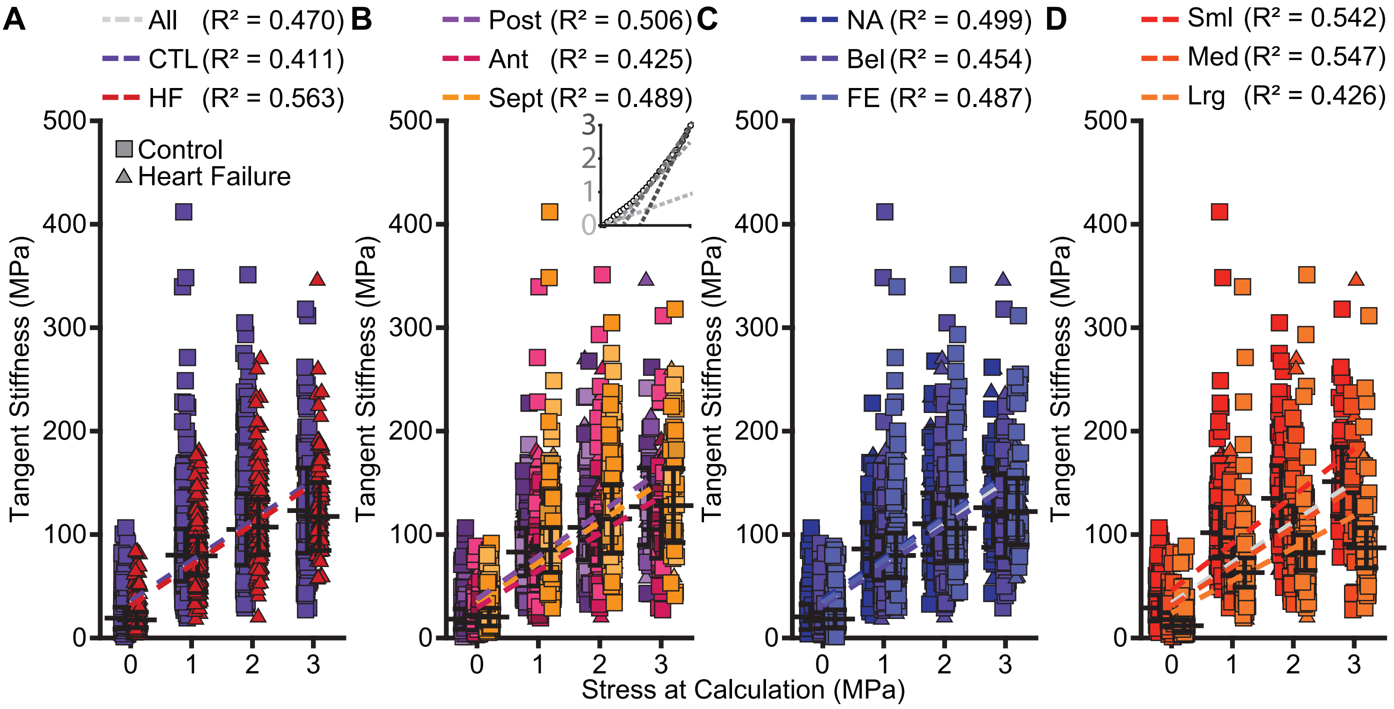


Figure S15: Tangent stiffness increases as a function of where the stiffness is taken on the stress-stretch curve. Specimens are grouped by **(A)** heart failure, **(B)** leaflet, **(C)** insertion site, or **(D)** size, and the tangent stiffness is computed as the tangent of the stress-stretch curve near 0, 1, 2, or 3 MPa (see inset). A linear model is fit to the tangent stiffness versus stress at calculation for all (All) specimens together and for each group, and adjusted $R^2$ values are listed for these fits.

the cycle. Median normalized hysteresis was 0.254 [0.202, 0.316]. In Figure S16, we show the normalized hysteresis for each test when grouped by heart failure, leaflet, and insertion site. Interestingly, we found that only insertion site was significant ($p = 0.003$), where belly chordae dissipated more energy than free edge chordae ($p = 0.002$). Specimen cross-sectional area ($p = 0.073$), heart failure ($p = 0.210$), leaflet ($p = 0.322$), and peak stress ($p = 0.950$) were not significant. Thus, cross-sectional area does not appear to dictate the viscoelastic properties of the chordae like it does the mechanical properties.

We found that toe stiffness, calf stiffness, and transition stretch were associated with the peak stress reached during testing. We show the toe stiffness, calf stiffness, and transition stretch as a function of the peak stress reached in Figure S17. Values from 1 MPa tests were lower than those from 2 and 3 MPa tests (all $p < 0.001$). However, interactions between peak stress and condition, leaflet, or insertion site were not significant (all $p > 0.1$). That is, the effect of progressive loading is similar across specimens regardless of heart failure, leaflet, and insertion site as shown by the lack of a trend between peak stress and leaflet-by-insertion site in Figure S17.

Specimens underwent 10 cycles at each sequential loading stage, so higher-stress tests also followed more cumulative loading. Their responses could therefore reflect additional conditioning, stiffening, any damage, slippage, or other history-dependent effects due to the repeated loading. Because force and strain references were reset at each stage, the observed differences could reflect both loading history and changes in reference configuration. Importantly, we included peak stress as a fixed effect to adjust group comparisons for systematic differences among testing stages, but did not establish the absence of group-specific order effects. Constitutive fits combined these stage-referenced responses. The effects of reference resetting, linear strain-time smoothing, and fitting-region adjustments were not independently quantified.

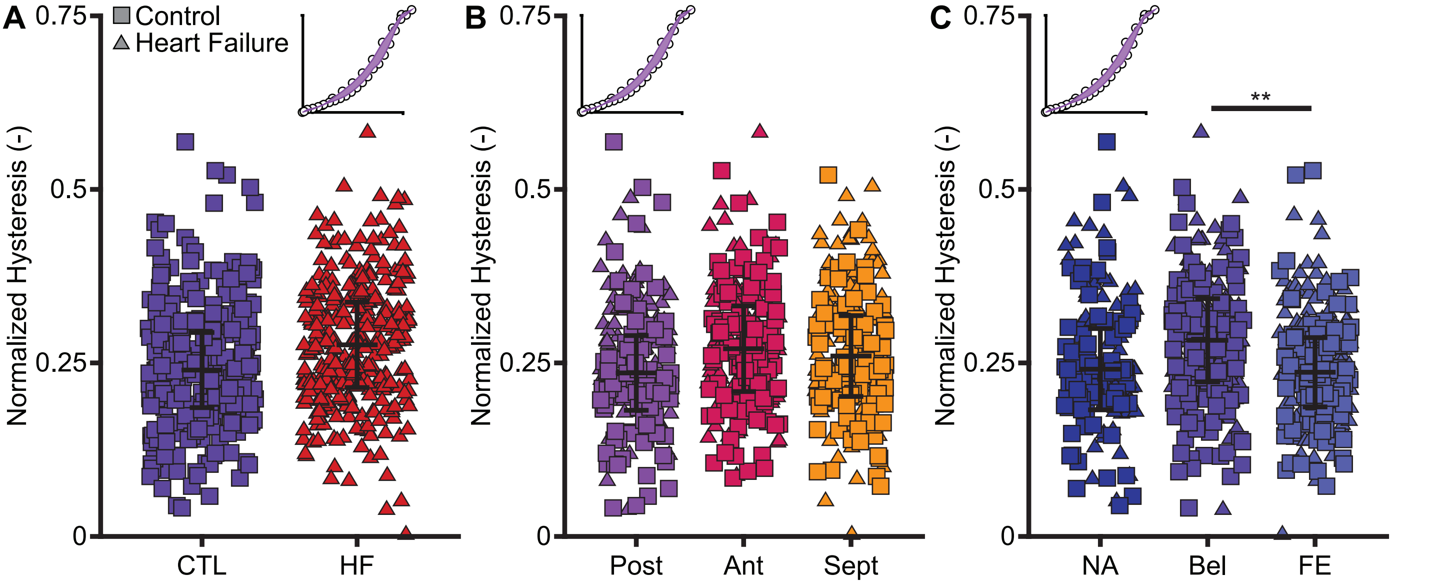


Figure S16: Chordae tendineae inserting into the belly dissipate relatively more energy, i.e., show greater normalized hysteresis than those inserting into the free edge. Specimens are grouped by **(A)** heart failure, **(B)** leaflet, or **(C)** insertion site after accounting for the cross-sectional area of each specimen, where each point is from one of 689 uniaxial tests, where error bars indicate the median and interquartile range. Normalized hysteresis is calculated as the difference between the loading and unloading energy densities (see insets) divided by the loading energy density. Only tests with a valid insertion site are shown. Asterisks indicate statistical significance (* p<0.05, ** p<0.01, *** p<0.001).

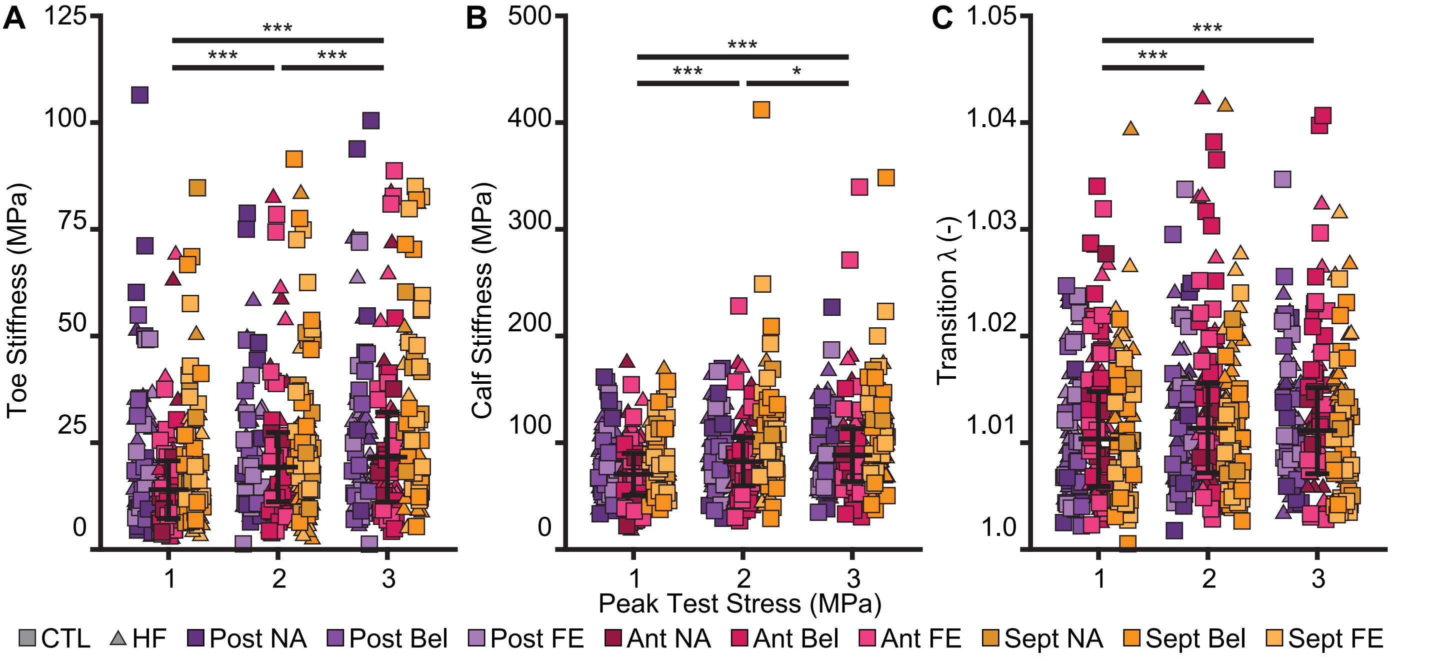


Figure S17: **(A)** Toe stiffness, **(B)** calf stiffness, and **(C)** transition stretch increase when the specimen is tested to a larger peak stress. That is, repeated loading systematically increases the computed mechanical quantities in a similar manner between groups. Each point is from one of 689 uniaxial tests, where error bars indicate the median and interquartile range. Only tests with a valid insertion site are shown. Asterisks indicate statistical significance (* p<0.05, ** p<0.01, *** p<0.001).
